# Spatiotemporal Dynamics of Vcam1 Regulates Cholangiocarcinoma Mass Expansion and Tumor Dissemination under Growth-suppressive Peritumoral Myofibroblasts

**DOI:** 10.1101/2023.01.24.525379

**Authors:** Cheng Tian, Liyuan Li, Qingfei Pan, Beisi Xu, Yizhen Li, Li Fan, Anthony Brown, Michelle Morrison, Kaushik Dey, Jun J. Yang, Jiyang Yu, Evan S Glazer, Liqin Zhu

## Abstract

Intrahepatic cholangiocarcinoma (iCCA) is characterized by its highly desmoplastic stroma. Myofibroblasts (MFs) are present both within the tumor mass (intratumoral MFs, iMFs) and at the tumor border (peritumoral MFs, pMFs). Using a spheroid-based coculture system, we show that the initial iCCA-pMF contact is growth suppressive to the tumor cells. However, prolonged iCCA-pMF interaction elicits significant tumor cell invasion and dissemination. We find that vascular cell adhesion molecule-1 (*Vcam1*) level is elevated in tumor cells in contact with pMFs but low in disseminated tumor cells both in vitro and in vivo. A gene regulatory network analysis of mouse and patient iCCA tumors and *Vcam1* knockout (*Vcam1*^*KO*^) demonstrate a heavy involvement of Vcam1 in epithelial-to-mesenchymal transition. While *Vcam1*^*KO*^ has only a limited impact on tumor cell growth in their monoculture, *Vcam1*^*KO*^ spheroids exhibit instant dissemination and a severe growth defect when cocultured with pMFs. When transplanted into the liver, *Vcam1*^*KO*^ iCCA cells show a similar increase in dissemination but a significant defect in establishing primary and metastatic tumors. Incomplete blocking of Vcam1 in vivo reduces the size but increase the number of metastatic lesions. Overall, our study shows a spatiotemporal regulation of iCCA growth and dissemination by pMFs in a Vcam1-dependent manner.

## INTRODUCTION

Cholangiocarcinoma is the second most common hepatic tumor after hepatocellular carcinoma (HCC). It is believed to originate from the biliary tracts within or outside the liver which lead to the development of intrahepatic or extrahepatic cholangiocarcinoma (iCCA or eCCA, respectively). iCCA has a worse prognosis than eCCA and is one of the deadliest cancers overall [1, 2]. One of the most prominent characteristics of iCCA is its highly desmoplastic stroma. This has led to many investigations on cancer-associated fibroblasts (CAFs) in iCCA development [3-7]. Indeed, the liver is one of the few internal organs that are developmental equipped with a complex fibrotic response machinery in order to protect its vital function in metabolism and immunity [8]. Nearly every chronic liver condition results in liver fibrosis eventually [9]. Hepatic stellate cells (HSCs), a major cell type giving rise to liver myofibroblasts (MFs) via their activation, have been demonstrated as the main source of iCCA CAFs [4, 10]. Therefore, it is conceivable that there is a spatiotemporal evolution of iCCA CAFs from the initial activation of HSCs at the tumor-liver border to their gradual incorporation into the tumor mass as tumor progresses.

Most iCCA CAF studies have focused on the MFs residing within iCCA tumor mass (intratumoral MFs, or iMFs) and found them being predominantly protumorigenic [3-7]. Contribution of MFs at the tumor border (peritumoral MFs, or pMFs) to iCCA development is less understood. Recent studies in HCC, although limited, found that pMFs are positively associated with HCC metastasis and recurrence [11, 12]. We reasoned that investigating iCCA pMFs was important to a more complete appreciation of the role of CAFs in this rare but highly aggressive cancer. To do so, we examined the distribution of both iMFs and pMFs in a cohort of early-stage iCCA patient tumors. We then tracked the spatiotemporal dynamics of the MFs in tumors collected from a metastatic iCCA orthotopic allograft model we derived from a *<u>P</u>rom1*^*CreERT2*^; *<u>P</u>ten*^*flx/flx*^; *<u>T</u>p53*^*flx/flx*^; *<u>R</u>osa-ZsGreen* (PPTR) liver cancer genetic model we established previously [13, 14]. We then focused on pMFs and assessed their impact on iCCA tumor cell growth and dissemination via multiple coculture systems we established in the laboratory. Lastly, we investigated the potential underlying molecular mechanisms that mediate the dynamic iCCA-pMF interaction.

## MATERIALS AND METHODS

### Mice

Two-month-old male and female C57BL/6J (B6) (Strain # 000664, The Jackson Laboratory, Bar Harbor, ME, USA) and *Crl:CD1-Foxn1*^*nu/nu*^ (CD-1 nude) mice (Strain Code 086, Charles River, Wilmington, MA, USA) were used for iCCA orthotopic transplantation and tail vein injection. *B6*.*129(Cg)-Gt(ROSA)26Sor*^*tm4(ACTB-tdTomato,-EGFP)Luo*^*/J* (Strain # 007676, The Jackson Laboratory, Bar Harbor, ME, USA) mice were used to isolated hepatocytes according to standard protocol. All mice were maintained in the Animal Resource Center at St. Jude Children’s Research Hospital. Animal protocols were approved by the St. Jude Animal Care and Use Committee.

### Reagents for cell labeling and detection

All flow cytometric analyses were performed in three biological replicates using a BD LSRFortessa™ cell analyzer (BD Biosciences) and flow data were analyzed using FlowJo_V10. See Supplementary Material for details. Vcam1 detection: APC anti-mouse CD106 (Vcam1) antibody (Cat. No. 105717, BioLegend, San Diego, CA, USA). Cell proliferation: CellTracker™ Red CMTPX dye (Cat. No. C34552, Invitrogen, Carlsbad, CA, USA).

### Cell Culture

PPTR cells were established from *<u>P</u>rom1*^*CreERT2*^; *<u>P</u>ten*^*flx/flx*^; *<u>T</u>p53*^*flx/flx*^; *<u>R</u>osa-ZsGreen* (PPTR) liver cancer organoids [14] and adapted to 2D culture.

### Cytokine array assay

Conditioned media (CM) were collected from cell cultures after four days. The levels of secreted cytokines in the CM were measured using Mouse Cytokine Antibody Array C3 (Cat. No. AAM-CYT-3-8, RayBiotech, GA, USA) according to the manufacturer’s instructions.

### Generation of *Vcam1*^*KO*^ PPTR cells via CRISPR/Cas9

*Vcam1*^*KO*^ PPTR cells were generated via a CRISPR/Cas9 approach. The sgRNA sequence 5′-AGACAGCCCACTAAACGCGANGG-3′ is located on the Exon 2 of the mouse *Vcam1* gene.

### Vcam1 neutralization

For in vivo assays, mice were treated three days after TVI transplantation with *InVivo*MAb anti-mouse CD106 (Vcam1) antibody (Vcam1^Ab^) (Cat. No. BE0027, Bioxcell, Lebanon, NH) or rat IgG (Cat. No. Mab006, R&D System, Minneapolis, MN) at 0.25 mg/mice via TVI for 3 weeks, twice a week, five mice/group. For in vitro assays, cultured cells were treated with 10 μM Vcam1^Ab^ or IgG for 4 days.

### Gene regulatory network analysis

A scalable software was used for gene regulatory network reverse-engineering from big data, SJARACNe (v-0.1.0) [15], to reconstruct context-dependent signaling interactomes of Vcam1.

### RNA sequencing and analysis

The mRNA libraries of *Vcam1*^*Ctrl*^ and *Vcam1*^*KO*^ PPTR cells collected from three different passages were constructed using Illumina TrueSeq stranded mRNA library prep kit and and paired-end 100-cycle sequencing was performed on Illumina NovaSeq sequencers per the manufacturer’s directions (Illumina).

### Study approval and patient samples

Animal experiments were approved by St. Jude Animal Care and Use Committee. The de-identified iCCA patient tumor samples were obtained under a protocol approved by the Institutional Review Boards at both St. Jude Children’s Research Hospital and The University of Tennessee Health Science Center.

### Data Availability Statement

The total mRNA sequencing data generated in this study will be publicly available. Gene Expression Omnibus (GEO) submission is pending.

Detailed methods can be found in the Supplementary Materials.

## RESULTS

### Accumulation of pMFs in iCCA patient and mouse allograft tumors

We performed IHC of alpha-smooth muscle actin (αSMA), a well-recognized MF marker, on Stage I iCCA patient tumors to examine their MF content. To avoid therapy-induced liver fibrosis, we selected three tumors resected with no neoadjuvant treatment and with >=5 mm tumor-surrounding liver attached. These tumors were limited due to the rareness and often late diagnosis of iCCA. We defined peritumoral as regions within 100 μm away from the tumor border from both the tumor and liver side, and tumor core as >=2mm from the border. In these tumors, we found αSMA^+^ iMFs in the tumor core and pMFs in the tumor-surrounding liver as expected (**Figure 1A)**. In particular, there was a consistently higher density of pMFs at the tumor border in these three tumors compared to the tumor core (**Figure 1B)**.

**Figure 1.**
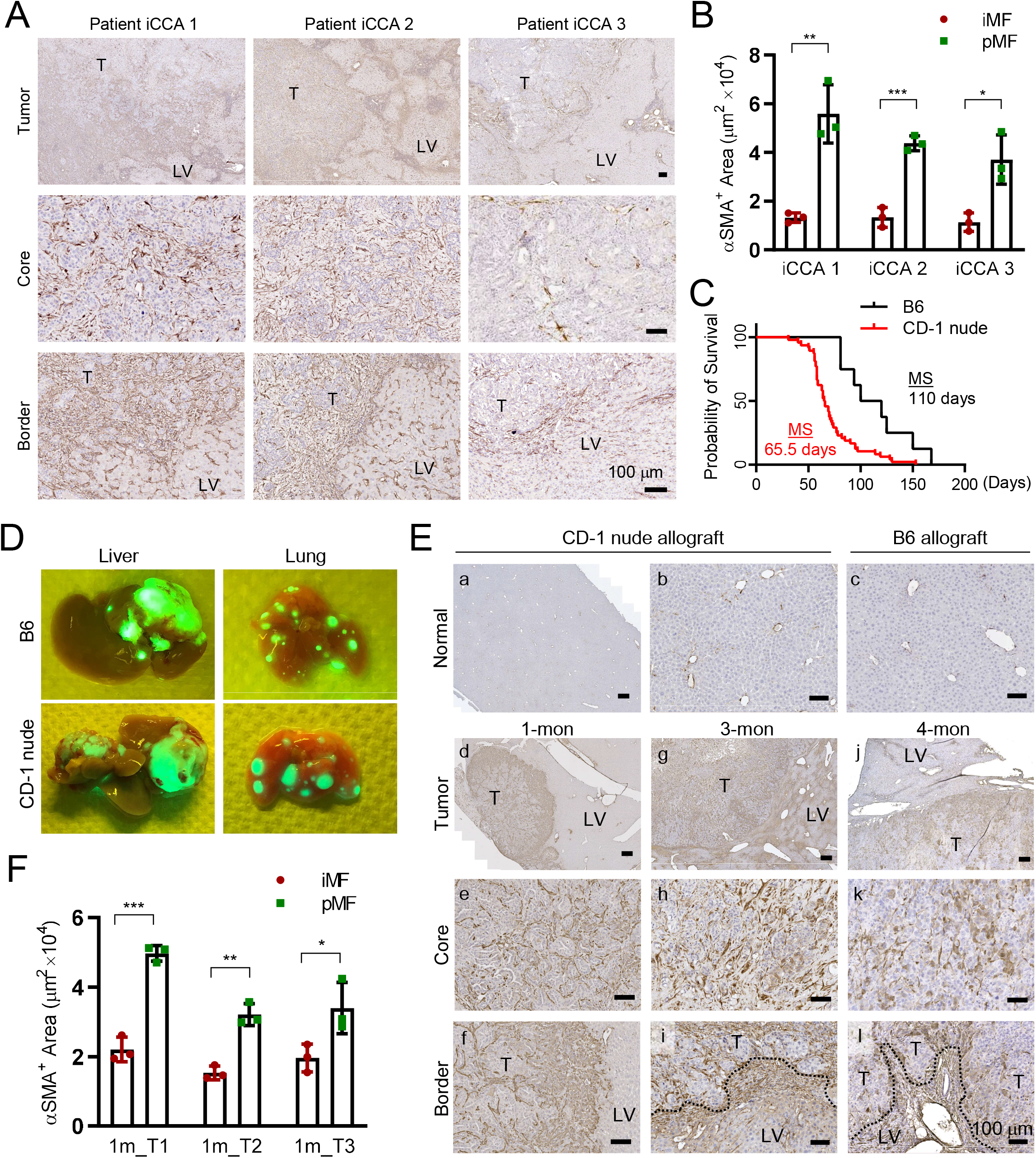
Accumulation of pMFs in mouse and patient iCCA tumors. **(A)** IHC of αSMA in three low-risk iCCA patient tumors showing higher density of MFs in the peritumoral region than the tumor core. **(B)** Quantitative comparison of the area occupied by the iMFs and pMFs in the three patient tumors in (A). Three regions with the heaviest iMF and pMF accumulation were measured in each tumor. **(C)** The animal survival curves of CD-1 nude and B6 mice orthotopic transplanted with PPTR tumor cells. **(D)** Gross GFP fluorescence images of the liver and lung from the iCCA allograft models showing the intrahepatic and lung metastases. **(E)** IHC of αSMA on wildtype liver (a-c) and those from the CD-1 nude (d-i) and B6 (j-l) allograft models at the indicated time points. Dotted lines: invasive tumor border; arrows: αSMA^+^ pMFs; T: tumor; LV: liver. All scale bars are 100 μm. **(F)** Quantitative comparison of the area occupied by the iMFs and pMFs in 1-mon iCCA allograft tumors. Three regions with the heaviest iMF and pMF accumulation were measured from three tumors.

Since MF activation in iCCA patients could be triggered either by tumorigenesis or by other non-tumor-related preexisting disease conditions, we utilized an orthotopic allograft model of metastatic iCCA we established in our laboratory to track tumor-induce MF activation. We have reported an allograft models of metastatic iCCA generated by orthotopically transplanting tumor cells cultured from a PPTR liver cancer genetic model we previously established [13, 14]. Metastatic iCCA tumors were generated in both CD-1 nude and B6 mice with tumor developing faster and more consistently in the former (**Figure 1C**). Tumors in both models are similar and recapitulate patient tumor histologically (**Supplemental Figure S1A**). The *Rosa-ZsGreen* (*ZsG*) reporter allele in the PPTR tumor cells enabled direct visualization of the tumor masses as well as disseminated tumor cells (DTCs) (**Figure 1D** and **Supplemental Figure S1B**). We collected the iCCA allograft tumors at different time points and performed αSMA IHC. In normal mouse liver, αSMA positivity was only found in the smooth muscle cells lining large vessels (**Figure 1E, a-c**). In the iCCA tumors collected after one month of transplantation from CD-1 nude mice, we noticed the accumulation of αSMA^+^ pMFs at the tumor border similar to patient tumors (**Figure 1E, d**), indicating MF activation in the host liver elicited by iCCA development. Alpha-SMA^+^ iMFs were also present in the tumor core, however, at a lower density than those in the peritumoral region (**Figure 1E, e & f**, and **Figure 1F**). In the 3-month tumors collected from the CD-1 nude allografts, there was an evident increase of αSMA^+^ iMFs within the tumor core although a spatial heterogeneity was observed (**Figure 1E, g-h**). Accumulation of pMFs at the tumor border also became more heterogeneous and was mostly found at the tumor border where invasive tumor clusters were emerging (**Figure 1E, i**). Four-month tumors collected at the end point from the B6 allograft model showed a similar presence of dense iMFs in the tumor core and pMF accumulation particularly at the invasive border (**Figure 1E, j-l**). These observations in the iCCA orthotopic allograft model suggest pMF accumulation on the tumor border as an early event in iCCA development and their potential contribution to tumor invasion at later stages. Because of the faster and more consistent tumor development in the CD-1 allograft model, we performed all following in vivo experiments in this background.

### Peritumoral MFs suppress, and intratumoral MFs promote, iCCA growth in vitro

Previous studies have consistently shown that CAFs within iCCA tumor mass promote tumor cell growth [3, 4, 16]. When examining tumor cell proliferation in our allograft tumors in association with the presence of pMFs and iMFs, we found tumor cells in the pMF^high^ regions in the 1-month allograft tumors had significantly fewer Ki67^+^ cells than those in the tumor core (**Figure 2A** and **2B**). In the 3-month tumors where iMF^high^ regions were emerging in the tumor core, we noticed a strong, positive association between the iMFs content and Ki67 positivity (**Figure 2A** and **2B**). However, at the invasive tumor border where pMFs accumulated, tumor cell proliferation rate was lower than that of the iMF^high^ tumor core (**Figure 2A** and **2B**). These results suggest that iMFs promote tumor cell proliferation as previously reported, but pMFs are potentially growth-suppressive.

**Figure 2.**
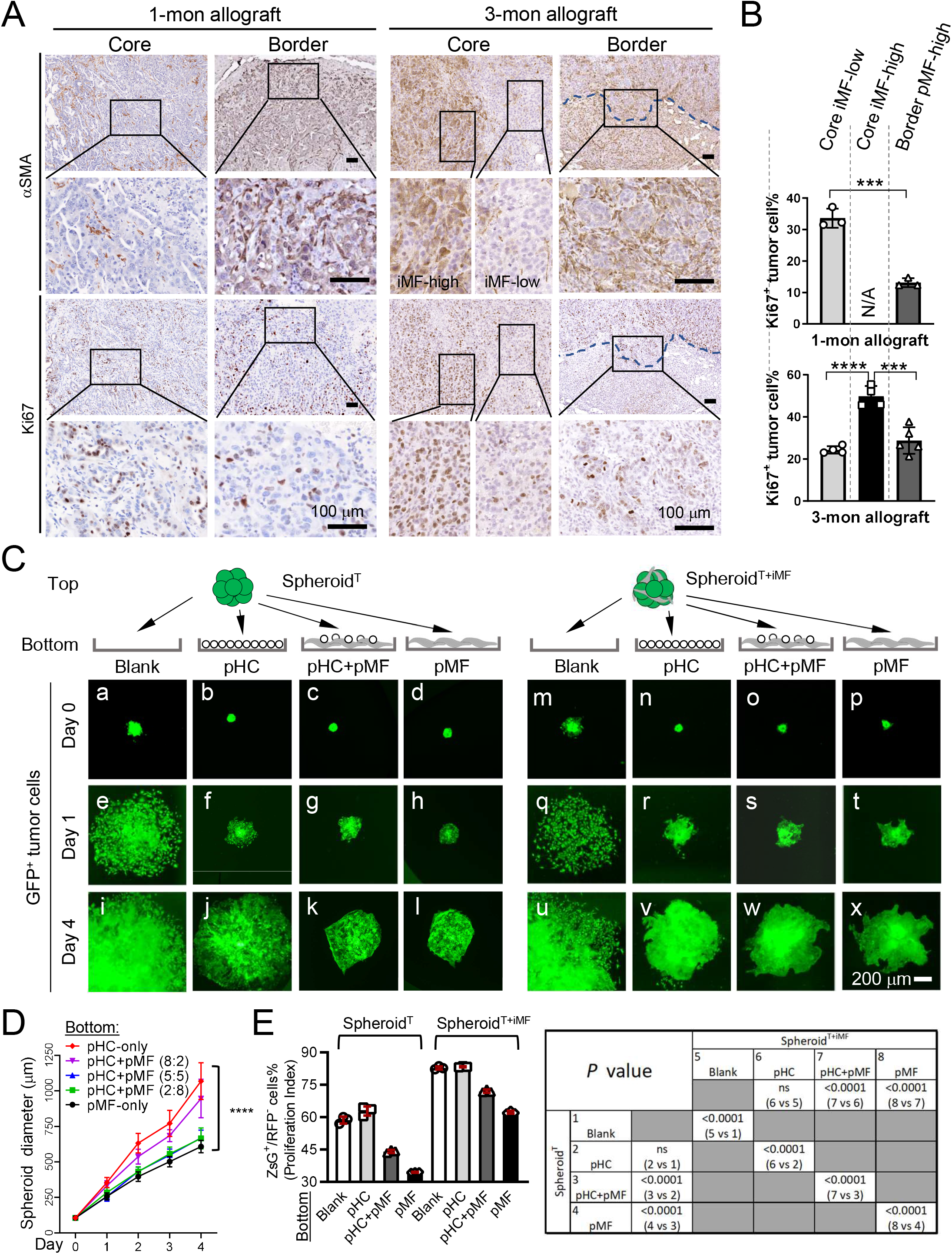
Peritumoral MFs suppress, and intratumoral MFs promote, iCCA growth in vitro. **(A)** Alpha-SMA and Ki67 IHC in the 1-month and 3-month iCCA allograft tumors. Images in the same row share the scale bar of 100 μm. **(B)** Quantification of Ki67^+^ cells in the indicated areas of the 1-and 3-month iCCA allograft tumors. N/A: No core iMF-high regions present in the 1-month tumors. Three tumors were examined at each time point. Student *t*-test, *P* value *** <0.001, ****<0.0001. **(C)** Time course images of the ZsG^+^ Spheroid^T^ and Spheroid^T+iMF^ in the 2.5D coculture with the indicated bottom layers. All images share the same 200 μm scale bar. **(D)** Quantification of the diameters of Spheroid^T^ in the indicated coculture conditions. Two-way ANOVA comparison, *P* value ****<0.0001. **(E)** Flow cytometry-based detection of the CellTracker-RFP dye of Spheroid^T^ and Spheroid^T+iMF^ placed on the indicated bottom layer. Note that RFP-negative (RFP^-^) cells are plotted as more RFP^-^ cells indicates higher tumor cell proliferation index. *P* values of the Student *t*-test between the indicated groups are shown on the right.

To assess the effect of pMFs on tumor cell proliferation directly, we developed a spheroid-based “2.5-dimensional” (2.5D) tumor-pMF coculture system by placing 3D tumor spheroids (Spheroid^T^) onto a 2D layer of pMFs (see the schematic illustration in **Figure 2C**). Since HSCs are the main source of liver MFs [4], we acquired primary mouse HSCs and induced their activation to MFs via 2D culture on plastic plates [17]. Their high levels of αSMA was confirmed via immunofluorescence after three passages in culture (**Supplemental Figure S2**). Freshly isolated primary mouse hepatocytes were also used in the 2D layer as a comparison to MFs. To assess tumor cell proliferation, PPTR tumor cells were labeled with CellTracker™ Red CMTPX dye (CellTracker-RFP). CellTracker-RFP is be diluted into daughter cells upon cell division. Therefore, a lower RFP intensity indicates a faster cell proliferation. CellTracker-RFP-labeled PPTR cells were used to generate Spheroid^T^, or mixed at 1:1 with the mouse HSC-derived MFs to generate spheroids with iMFs (Spheroid^T+iMF^). Four types of 2D bottom layer were used in the cocultures: control blank (no bottom 2D layer), freshly isolated hepatocytes (peritumoral hepatocytes, or pHC), a 1:1 mixture of hepatocytes and MFs (pHC+pMF), and MFs only (pMF) (**Figure 2C**).

Cells in the bottom layer were seeded at a same total number and allowed to attach for 24 hr before Spheroid^T^ or Spheroid^T+iMF^ were seeded on top. Tumor spheroids gradually flatted down within 24 hr and a slower expansion of the spheroid ZsG area was noticed in the wells with a bottom layer than those without (**Figure 2C, f-h vs. e**). Compared to pHCs, pMFs exhibited a stronger suppression on tumor spheroid expansion (**Figure 2C, l vs. j**). To confirm, we also placed Spheroid^T^ on the top of pHC+pMF mixed at different ratios and found a significant and inversed association between Spheroid^T^ area and the pMF content (**Figure 2D**). The similar suppressive effect of pMFs was also observed in the cocultures with Spheroid^T+iMF^ although at a lesser extent compared to Spheroid^T^ (**Figure 2C, m-x**). Tumor cell proliferation in the cocultures were examined after four days by measuring the CellTracker-RFP intensity in ZsG^+^ tumor cells. In both Spheroid^T^ and Spheroid^T+iMF^ cocultures, we found higher numbers of RFP^+^ cells as well as higher RFP intensity in the conditions with higher pMF contents, indicating a slower tumor cell growth with more pMFs in the bottom layer (**Figure 2E**, and **Supplemental Figure S3**). When comparing Spheroid^T^ and Spheroid^T+iMF^ placed on the same bottom layer, tumor cells in Spheroid^T+iMF^ showed a consistently and significantly higher proliferation rate indicated by their lower levels of CellTracker-RFP (**Figure 2E**, and **Supplemental Figure S3**), consistent with the previous findings that iCCA CAFs within the tumor mass were pro-proliferative [4, 18]. Overall, these results are consistent with our in vivo observations in the iCCA allograft tumors, that iMFs and pMFs affect iCCA growth differently – iMFs promote and pMFs suppress tumor cell proliferation.

### Elongated iCCA-pMF interaction elicits iCCA cell dissemination in vitro

To study the long-term effect of pMFs on tumor cell behaviors, we set up long-term 2.5D cocultures similar to Figure 2 and monitored tumor spheroids up to six weeks. We noticed that Spheroid^T^ placed on top of pMF continued to grow more slowly than those on blank and pHC (**Figure 3A, a-d and i-l**). We then started noticed tumor cell invasion only in the conditions with pMFs which became evident by two weeks (**Figure 3A, f & h**). When Spheroid^T^ were placed on pHC+pMF mixed at different ratio, there was an association between higher pMF content and longer invasive processes (**Figure 3B**). Tumor cell dissemination was also more evident when pMFs were present (**Figure 3A, h, inset**). Compared to Spheroid^T^, Spheroid^T+iMF^ appeared to have a more invasive tumor border on the two pMF-containing conditions although, interestingly, no evident tumor cell dissemination was observed (**Figure 3A, i-p**).

**Figure 3.**
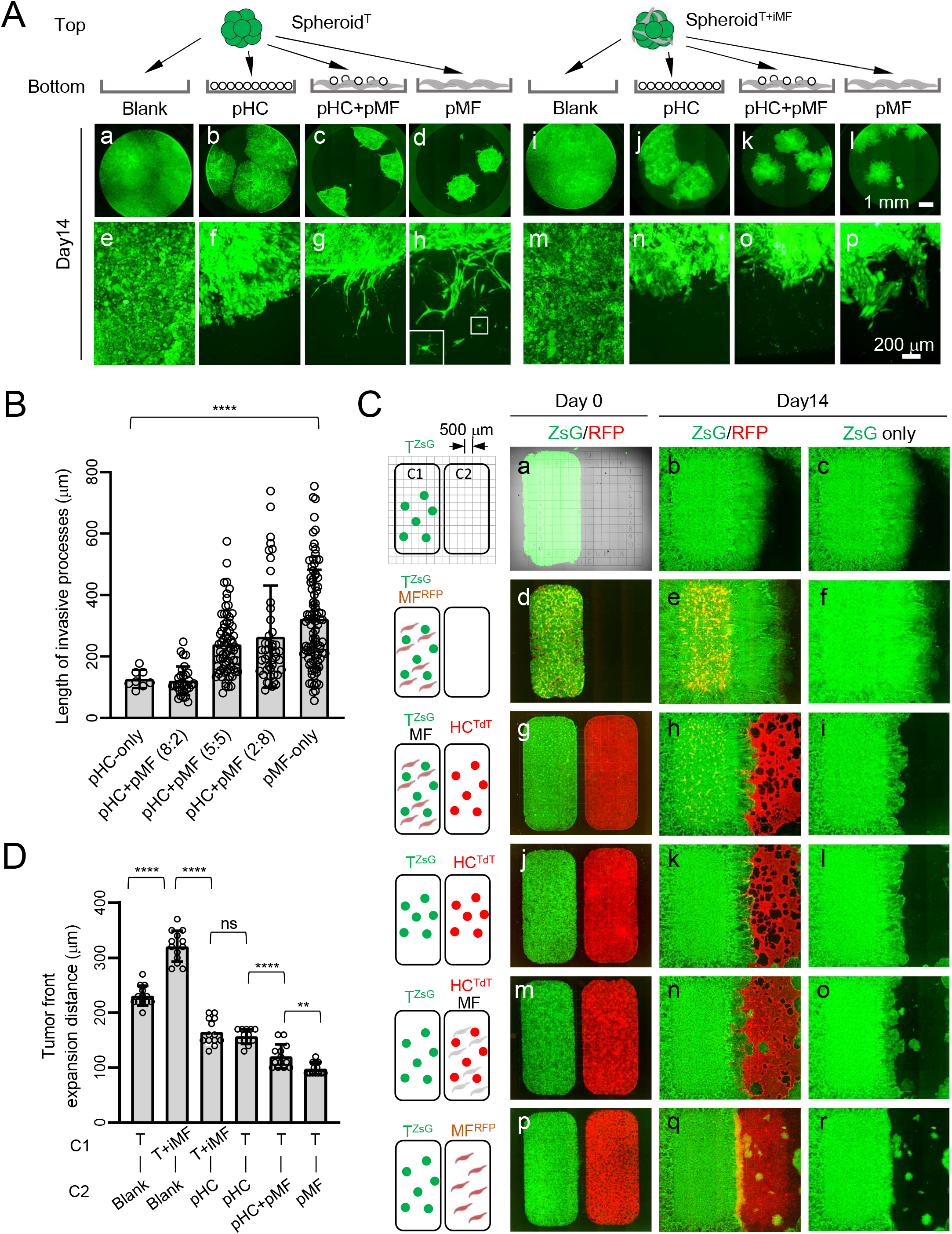
Elongated iCCA-pMF interaction induces tumor cell dissemination. **(A)** ZsG fluorescence images of Day 14 Spheroid^T^ and Spheroid^T+iMF^ 2.5D cocultures on the indicated bottom layers. Images on the same row share the same scale bar. Inset in **h**: higher magnification image of the boxed disseminated tumor cell. **(B)** Day 0 and 14 fluorescence images of the indicated 2D two-chamber cocultures. MFs are labeled with CellTracker-RFP dye in the conditions when no TdT^+^ HCs were added (**d-f** and **p-r**). Clonal tumor dissemination only occurred when pMFs were present in the C2 chamber (**m-r**). **(C)** Measurement of the expanding distance of the right tumor border in the indicated conditions. Student *t*-test, *P* value: ns, not significant, ** <0.01, ****<0.0001. **(D)** Day 1 and Day 14 images of the 2.5D coculture of human iCCA cell line HuCCT1 and human activated HSC cell line LX2. HuCCT1 cells were GFP^+^. Arrows in (**j, l**): disseminated HuCCT1 tumor spheroids when LX2 cells were placed outside of tumor spheroids as pMFs.

To better visualize the effect of pMFs and iMFs on tumor cell invasion and dissemination, we developed a 2D coculture system using a two-chamber culture insert to spatially separate tumor cells and MFs (C1: left chamber; C2: right chamber) (**Figure 3C**). Culture slides engraved with a 500 μm-grid were used to measure tumor area expansion. Different combinations of hepatocytes, MFs, and PPTR tumor cells were seeded as indicated in **Figure 3C**. The inserts were removed after 24 hours to allow cells from the two chambers to interact. We similarly found that iMFs promoted iCCA cell growth in this two-chamber cocultures indicated by a faster expansion of the tumor area (**Figure 3C, a-c vs. d-f**, and **Figure 3D**). Placing pHCs in C2 slowed down tumor expansion and reduced the growth-promoting effect of iMFs (**Figure 3C, g-i vs. j-l** and **Figure 3D**). Tumor cell expansion rate was further reduced when pMFs were mixed with pHCs in C2 which was accompanied by a marked increase in tumor cell invasion and dissemination by two weeks (**Figure 3C, j-l vs. m-o** and **Figure 3D**). When only pMFs were placed in C2, tumor cells expanded the slowest while disseminating the most (**Figure 3C, p-r**, and **Figure 3D**). To further validate the effect of pMFs and iMFs on human liver cancer cells, we tested the interaction between a human hepatoma cell line HepG2 and a human activated HSC cell line LX2 [19] since there were no commercially available human iCCA cell lines. In the same 2.5D cocultures, we found that pLX2 similarly suppressed tumor spheroid growth but induced evident tumor cell dissemination. In contrast, iLX2 significantly increased HepG2 spheroid growth without clear induction of tumor cell dissemination (**Supplemental Figure S4**). Taken together, these results indicate that pMFs and iMFs play different role in promoting the aggressive behaviors of tumor cells, that pMFs retard tumor growth but elicit tumor dissemination while iMFs promote tumor growth with a lesser effect on tumor cell dissemination.

### iCCA-pMF interaction upregulates Vcam1 in tumor cells

Since the molecular mechanisms underlying iCCA-iMFs interaction have been investigated in the past by multiple groups [4, 18], we focused on pMFs in this study, in particular, how tumor cells responded to pMF suppression at the molecular level to explain our observations above. We turned to cytokines as they were known key players in MF-tumor interaction [20, 21]. We chose a large cytokine array pre-dotted with antibodies to 62 mouse cytokines (RayBiotech Mouse Cytokine Array C3) and blotted with conditioned media (CM) collected from Day 4 Spheroid^T^ 2.5D cocultures as indicated in **Figure 4A**. Among the 62 cytokines, Vascular Cell Adhesion Molecule 1 (Vcam1) showed a mild but significant increase in the T:pMF CM compared to the others (**Figure 4A, a-e**, and **Supplemental Figure S5**). *Vcam1* upregulation in the T:pMF coculture was confirmed by quantitative RT-PCR (**Figure 4A, f**). Flow cytometry also detected a higher number of Vcam1^+^ tumor cells in the T:pMF coculture compared to T-only culture on Day 4 (**Figure 4B**). Vcam1 IHC on 1-month PPTR allograft tumors confirmed that Vcam1^+^ cells were predominantly the tumor cells close to the border where pMFs accumulated (**Figure 4C** and **Supplemental Figure S6**). *VCAM1* expression was more heterogeneous in iCCA patient tumors, but was also found predominantly in tumor cells close to the tumor border with pMFs accumulation (**Figure 4D**).

**Figure 4.**
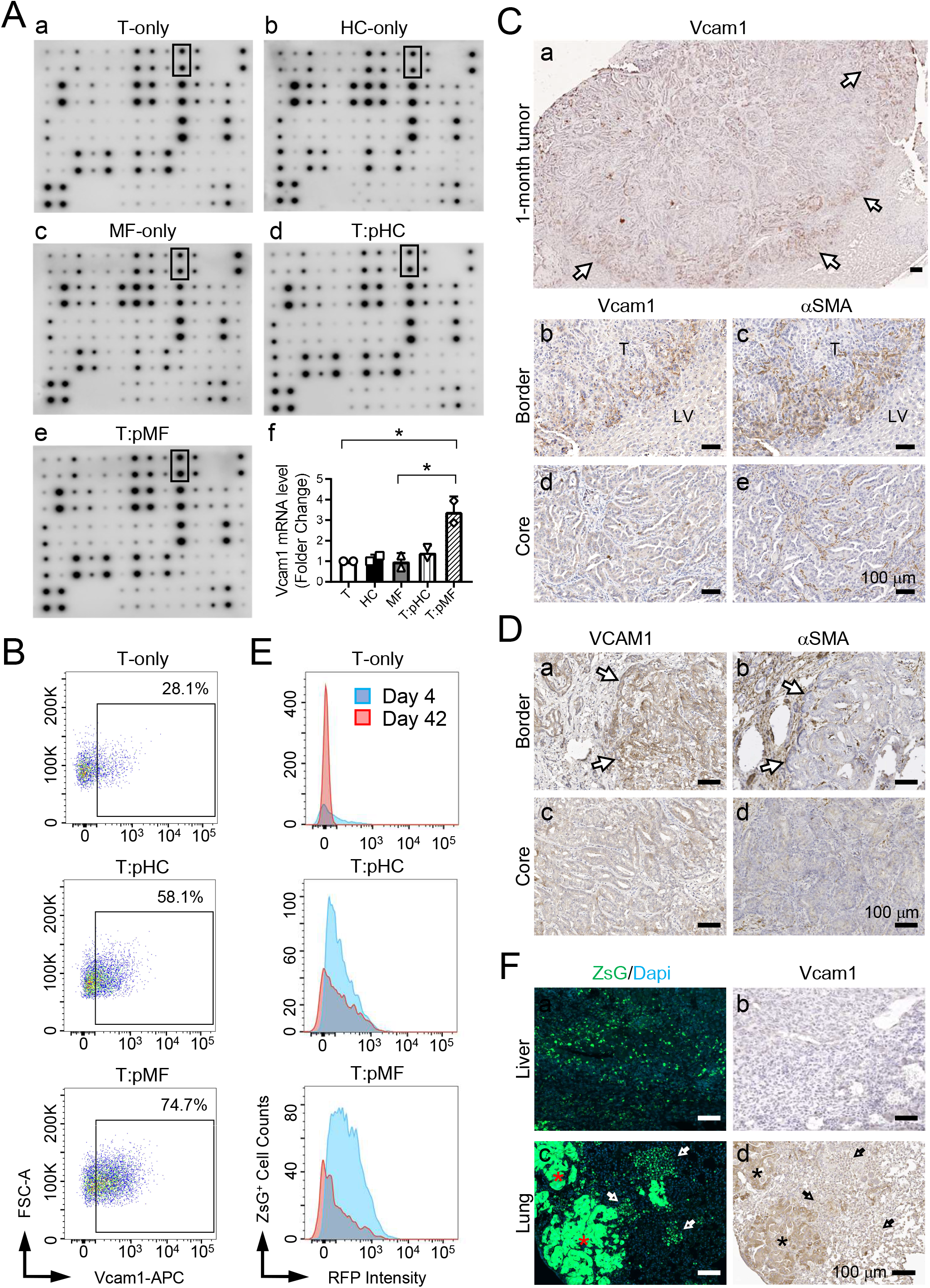
iCCA-pMF interaction induces *Vcam1* expression in the tumor cells. **(A)** Cytokine array assays using the CM from the indicated cultures on Day 4 (a-e). (**f**): Quantitative RT-PCR of *Vcam1* mRNA levels in the indicated cultures. Student *t*-test, *P* value * <0.05. **(B)** Vcam1 detection in the indicated 2.5D cocultures on Day 4 via flow cytometry. The percentage of Vcam1^+^ tumor cells in each condition was indicated. T:pMF coculture had the most Vcam1^+^ tumor cells. **(C)** Histogram of Vcam1 level in the tumor cells from the indicated cocultures on Day 4 and 42. There was a decrease in Vcam1 levels in Day 42 cultures. **(D)** Vcam1 and αSMA IHC on the serial sections of the indicated allograft tumors. All scale bars are 100 m. Arrows in **(a)**: Vcam1^high^ cells on the tumor border. Dotted lines in (**f, g**): tumor-pMF interface; arrows in (**h, i**): red arrows, Vcam1^+^ tumor cells next to iMFs; black arrows: Vcam1^-^ cells next to iMFs. **(E)** VCAM1 and αSMA IHC on serial sections of an iCCA patient tumor. Arrows: VCAM1^+^ tumor cells at the tumor border next to aSMA^+^ pMFs. **(F)** ZsG/Dapi fluorescence and Vcam1 IHC on the serial sections of the live (**a, b**) and lung (**c, d**) from iCCA allografts showing decreased Vcam1^high^ levels in the DTCs. Asterisks in (**c, d**): Vcam1^low^ tumor cells within the lung metastases; arrows in (**c, d**): Vcam1^low^ DTCs.

Vcam1 has been previously reported to promote solid tumor metastasis [22, 23]. However, interestingly, we found that Vcam1 levels dropped significantly in tumor cells collected from the 6-week Spheroid^T^:pMF 2.5D coculture (**Figure 4E**). Since many tumor cells had detached from Spheroid^T^ at this time (**Supplemental Figure S7**), we suspected that this reduction in Vcam1 might be associated with tumor cell dissemination. We found that, indeed, the ZsG^+^ DTCs in both liver and lung in 3-month allograft tumors were consistently Vcam1-negative while tumor cells in the overt lung metastases had high levels of Vcam1 (**Figure 4F**). These observations suggest that Vcam1 is likely critical for tumor mass expansion in a growth-suppressive peritumoral microenvironment but a negative regulator of tumor cell dissemination.

### Vcam1 is involved in epithelial-to mesenchymal transition pathway in both mouse and human iCCA tumors

Our data above suggested that Vcam1 might be involved in epithelial-to-mesenchymal transition (EMT). To test this hypothesis, we first utilized a system biology approach to unbiased identify Vcam1 regulatory pathways in both human and mouse iCCA tumors. We have previously performed a large RNA-seq transcriptomic profiling of the tumors from the PPTR genetic and orthotopic transplantation models (N=46) [14]. We also identified 36 iCCA patient tumor samples from TCGA (The Cancer Genome Atlas)-CHOL (cholangiocarcinoma) database (**Figure 5A**). We then used a scalable software for gene regulatory network reverse-engineering from big data, SJARACNe (v-0.1.0) [15], to reconstruct the context-dependent signaling interactomes of Vcam1 using these two datasets. We found that, indeed, EMT pathway was one of the top pathways predicted to be significantly involved in Vcam1 regulation in both mouse and patient tumors (**Figure 5B**). We then generated *Vcam1*^*KO*^ PPTR cells via CRISPR/Cas9 (**Supplemental Figure S8**). Two *Vcam1*^*KO*^ single-cell clones were generated (*Vcam1*^*KO1*^ and *Vcam1*^*KO2*^) and their loss of Vcam1 expression was confirmed by qRT-PCR and flow cytometry (**Figure 5C, D**).

**Figure 5.**
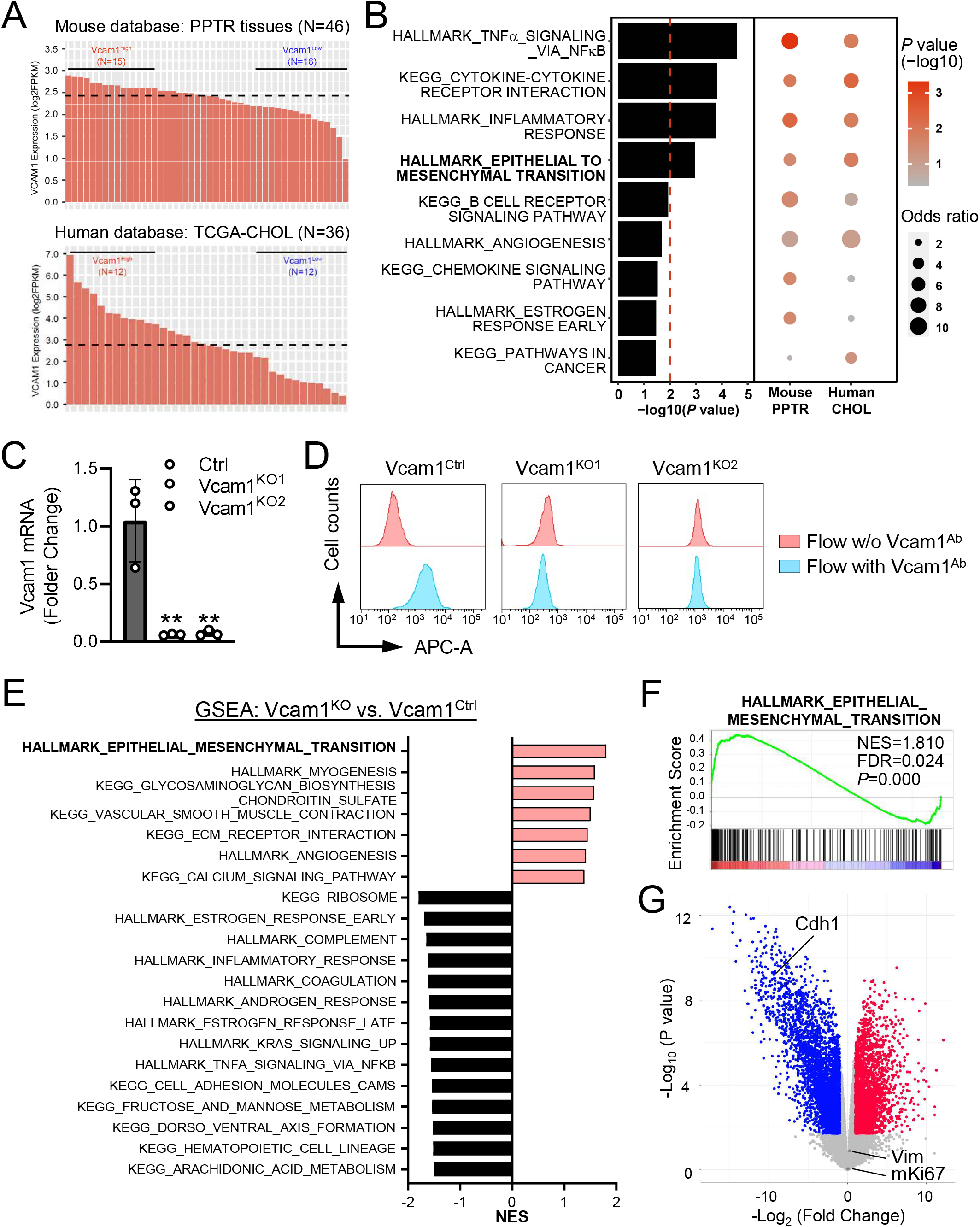
Vcam1 is involved in epithelial-to-mesenchymal transition in mouse and human iCCA tumors. **(A)** The mouse and human iCCA RNAseq datasets used in the analysis in (**B**). **(B)** Bubble plot representing gene set enrichment analysis (GESA) of predicted Vcam1 regulatory pathways in both mouse PPTR samples and human TCGA patient samples. Color intensity of the bubbles indicates the statistical significance of enrichment, and bubble size indicates the odds ratio. The bar plot on the left of the bubble plot shows the combined *P* values of the mouse and human datasets with the Stouffer method. Only the pathways with a combined *P* value < 0.05 are shown. **(C)** Quantitative RT-PCR of the *Vcam1*^*Ctrl*^ PPTR cells and two *Vcam1*^*KO*^ single-cell clones. Student *t* test. **, P < 0.01. **(D)** Flow cytometry of the *Vcam1*^*Ctrl*^ and *Vcam1*^*KO*^ cells. **(E)** GSEA analysis of the RNA-seq profiles of *Vcam1*^*KO*^ (*Vcam1*^*KO1*^ and *Vcam1*^*KO2*^ combined) vs. *Vcam1*^*Ctrl*^ cells. Pathways in red are upregulated in *Vcam1*^*KO*^ cells and pathways in black are downregulated. **(F)** The GSEA plot of HALLMARK_EPITHELIAL_MESENCHYMAL_TRANSITION pathway in *Vcam1*^*KO*^ vs. *Vcam1*^*Ctrl*^ cells. **(G)** The volcano plot of the gene expression fold change in *Vcam1*^*KO*^ vs. *Vcam1*^*Ctrl*^ cells.

RNA-seq transcriptomic profiling and gene set enrichment analysis (GSEA) confirmed that EMT pathway is the most significantly enriched pathway in the *Vcam1*^*KO*^ compared to the control (*Vcam1*^*ctrl*^) cells (**Figure 5E, F**). *Cdh1* gene (encoding E-cadherin), one of the marker genes associated with the epithelial features of cancer cells [24], was among the most downregulated genes in the *Vcam1*^*KO*^ cells (**Figure 5G**). No increase in *Vim* expression (encoding vimentin), a mesenchymal marker gene, was found in the *Vcam1*^*KO*^ cells likely because *Vcam1*^*ctrl*^ cells already expressed high levels of *Vim* [14]. No changes in the *Ki67* level (encoded by *mKi67* gene) were found in the *Vcam1*^*KO*^ cells (**Figure 5G**) which is consistent with our findings in the PPTR allograft tumors (**Supplemental Figure S9**) and as previously reported [25]. These data suggest an interesting decoupling of cell proliferation and Vcam1-dependent EMT in iCCA cells.

### Vcam1 depletion abolishes the growth of iCCA tumor cells under pMF suppression

To further unravel the relationship between iCCA cell growth and dissemination, pMFs, and Vcam1, we examined the growth of *Vcam1*^*ctrl*^ and *Vcam1*^*KO*^ cells in 2D monoculture, 3D spheroid monoculture, and spheroid-pMF coculture. In the 2D monocultures, we found both *Vcam1*^*KO*^ clones were able to expand steadily to reach full confluency although at a slowed rate compared to *Vcam1*^*ctrl*^ cells (**Figure 6A**). But while *Vcam1*^*ctrl*^ cells continued to doubling slowly afterwards, *Vcam1*^*KO*^ cells stopped growing and eventually died off (**Figure 6B, C**). In their 3D spheroid monoculture, *Vcam1*^*KO*^ cells were able to form spheroids in AggreWell (**Supplemental Figure S10**) and reach confluence when cultured alone (**Figure 6D**). However, when seeded on the top of pMFs, *Vcam1*^*KO*^ spheroids disseminated instantly and showed a severe growth defect afterwards compared to *Vcam1*^*ctrl*^ spheroids (**Figure 6E**). We noticed a reduction in their growth area by 14 days when the spheroids cultured without pMFs continued to grow (**Figure 6F**).

**Figure 6.**
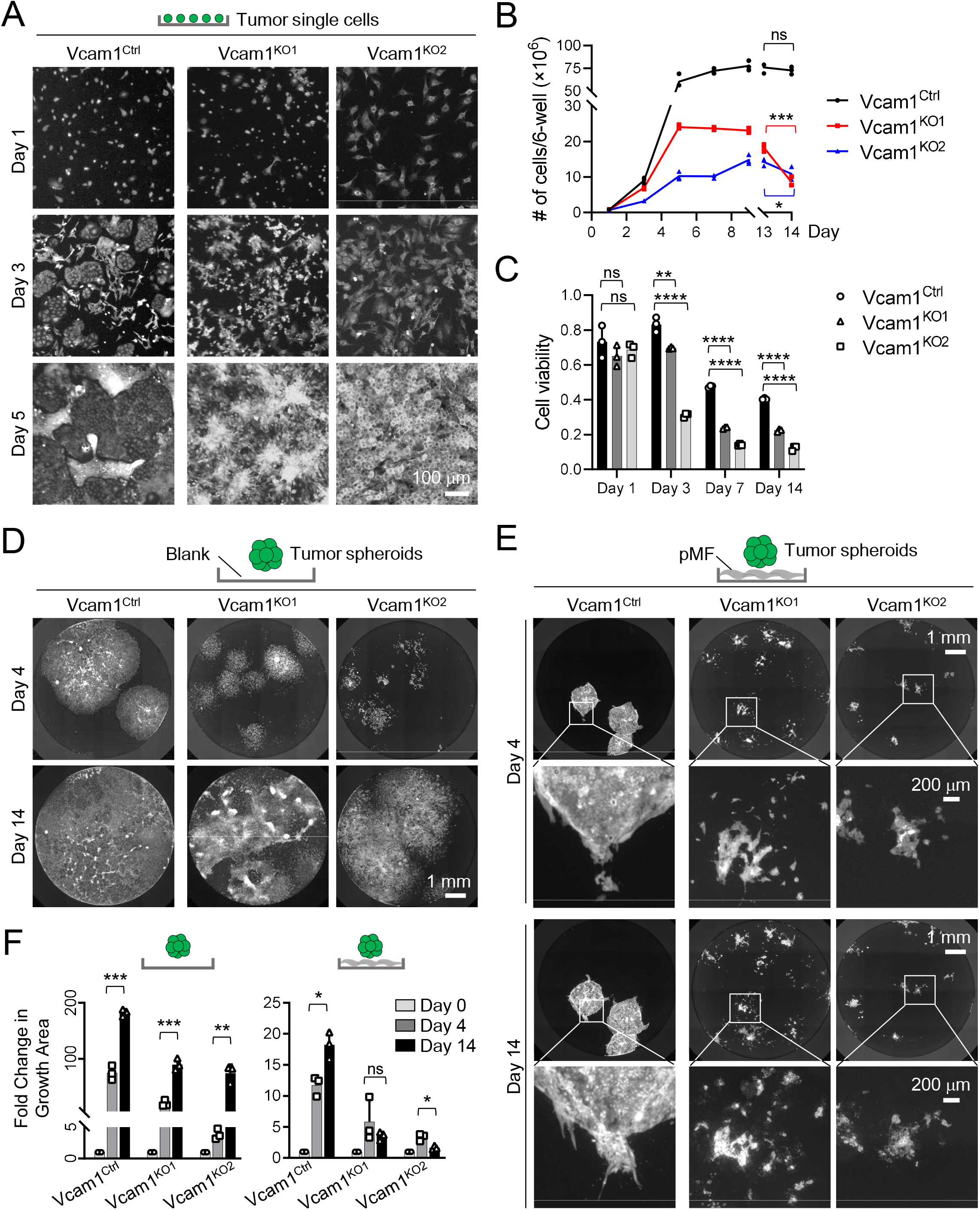
Vcam1 depletion severely impairs iCCA cell growth under pMF suppression. **(A)** GFP fluorescence images of the 2D monocultures of *Vcam1*^*Ctrl*^ and *Vcam1*^*KO*^ cells on Day 1, 3 and 5. **(B)** Cell number counts of *Vcam1*^*Ctrl*^ and *Vcam1*^*KO*^ cells in (A). Student *t*-test, *P* value, *** < 0.001, * < 0.05. **(C)** Quantification of the viable *Vcam1*^*Ctrl*^ and *Vcam1*^*KO*^ cells cultured in (A). **(D)** GFP fluorescence images of the spheroid monocultures of the *Vcam1*^*Ctrl*^ and *Vcam1*^*KO*^ PPTR cells on Day 4 and 14. **(E)** GFP fluorescence images of the tumor-pMF cocultures of the *Vcam1*^*Ctrl*^ and *Vcam1*^*KO*^ PPTR cells on Day 4 and 14. **(F)** Quantitative comparison of the growth area of the tumor cell spheroids in their mono-and co-culture. Statistics in (B), (C) & (F): Student *t*-test. *P* value, **** < 0.0001; ***< 0.001, ** < 0.01; * < 0.05.

When transplanted into the liver to allowed to grow for three weeks, *Vcam1*^*KO1*^ cells were less capable of establishing primary tumors compare to the control cells and failed to develop overt lung metastases (**Figure 7A**) (n=4 per group). However, gross GFP examination found a significantly higher number of individual DTCs in the lungs of all the animals transplanted with *Vcam1*^*KO1*^ cells than those with the control cells (**Figure 7B**). These results suggest that *Vcam1*^*KO*^ cells disseminate more readily than control cells but have an impaired ability to establish tumor mass. *Vcam1*^*KO2*^ clone was not tumorigenic in vivo (data not shown). We also tested the impact of Vcam1 loss on tumor growth in a tail vein injection (TVI) transplantation model. Although no liver pMF-tumor interaction was involved in this model, we reasoned that it was still a valid model for assessing Vcam1 function in iCCA development because of the high level of *Vcam1* expression in PPTR lung metastases (**Figure 4F**). Three weeks post injection, *Vcam1*^*KO1*^ cells showed a nearly complete failure in growing metastases while the control cells developed numerous tumors in the lung as expected. (**Figure 7C, D**) (n=4 per group). IHC of Vcam1, E-cadherin, and vimentin on *Vcam1*^*ctrl*^ and *Vcam1*^*KO*^ lung metastases confirmed a nearly completely loss of E-cadherin in the *Vcam1*^*KO*^ tumors without a clear difference in their vimentin levels (**Figure 7E, a-f**). Also consistent with the RNA-seq result, no differences in Ki67 positivity were observed between these two types of tumors (**Figure 7E, g vs. h**). Taken together, these results reveal an intereting context-dependent and Vcam1-regulated growth and dissemination of iCCA tumor cells. Vcam1 is not direcntly invovled in cell proliferation but support iCCA tumor mass expansion under pMF suppression and restrain tumor cell dissemination.

**Figure 7.**
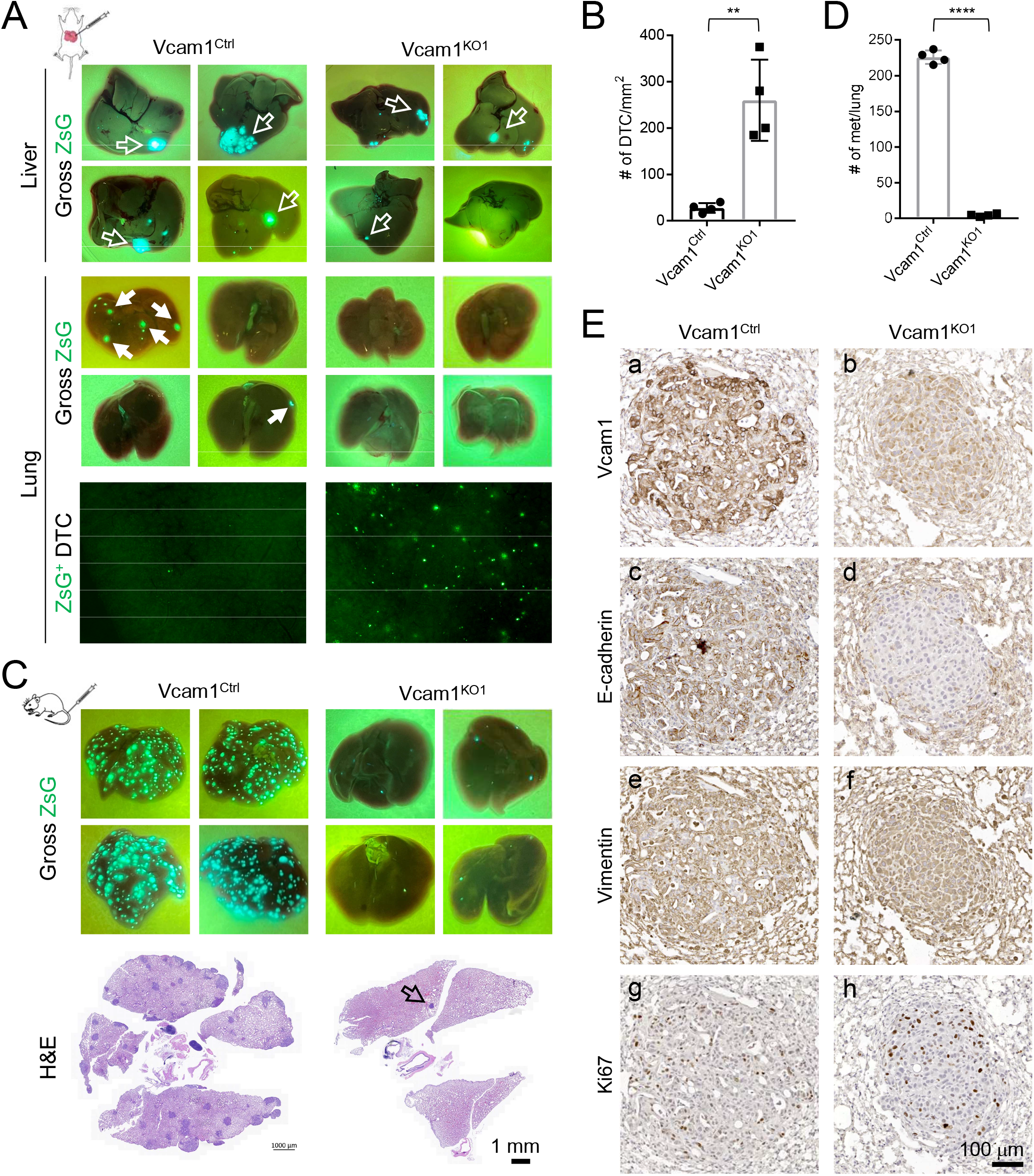
Vcam1 depletion abolished the metastatic growth of iCCA cells in vivo but promote the dissemination of individual tumor cells. **(A)** Top four rows: gross GFP images of the livers and lungs from mice orthotopically transplanted with *Vcam1*^*Ctrl*^ or *Vcam1*^*KO1*^ cells; bottom row: microscopic GFP images of the lungs from the *Vcam1*^*Ctrl*^ or *Vcam1*^*KO1*^ groups showing the higher number of DTCs in the latter. **(B)** Quantification of the numbers of DTCs in the lungs in (A). **(C)** Top two rows: gross GFP images of the lungs from TVI mice injected with *Vcam1*^*Ctrl*^ or *Vcam1*^*KO1*^ cells; bottom row: representative H&E images of the lungs from the *Vcam1*^*Ctrl*^ and *Vcam1*^*KO1*^ groups. **(D)** Quantification of the numbers of the lung metastases in (C). **(E)** The indicated IHC on serial sections of the *Vcam1*^*Ctrl*^ and *Vcam1*^*KO1*^ lung metastases. Note the complete loss of E-cadherin in the *Vcam1*^*KO*^ tumor (**d**). Statistics in (B) & (D): Student *t*-test. *P* value, **** < 0.0001; ** < 0.01.

### Partial neutralization of Vcam1 activity promotes iCCA metastasis in vivo

Finally, we tested the effect of a Vcam1 neutralizing antibody (Vcam1^Ab^) [26] on PPTR cell growth and dissemination in vitro and in vivo. Two PPTR lines were treated with Vcam1^Ab^ and they both showed increased migration ability (**Figure 8A**). Again, no differences in cell proliferation in their 2D monoculture were observed (**Figure 8B**). Since we found it challenging to statistically compare tumor growth in the orthotopic model due to the limited number of tumor masses and variation in size (**Figure 7A**), we performed the in vivo Vcam1^Ab^ treatment in the PPTR TVI model. We found the TVI group treated with Vcam1^Ab^ for three weeks developed more lung metastases than those treated with IgG (n=4 per group) (**Figure 8C**). However, metastases in the Vcam1^Ab^ group were smaller than those in the IgG group (**Figure 8D-F**). IHC showed that lower but only completely abolished Vcam1 levels in the tumors from the Vcam1^Ab^ group (**Figure 8G**). In both groups, DTCs showed no Vcam1 positivity, aSMA^+^ fibroblasts were commonly found around the Vcam1^+^ tumor cells (**Figure 8G**). These findings indicate that partial Vcam1 neutralization can be detrimental because it increases tumor cell dissemination but unable to completely prevent tumor cell growth.

**Figure 8.**
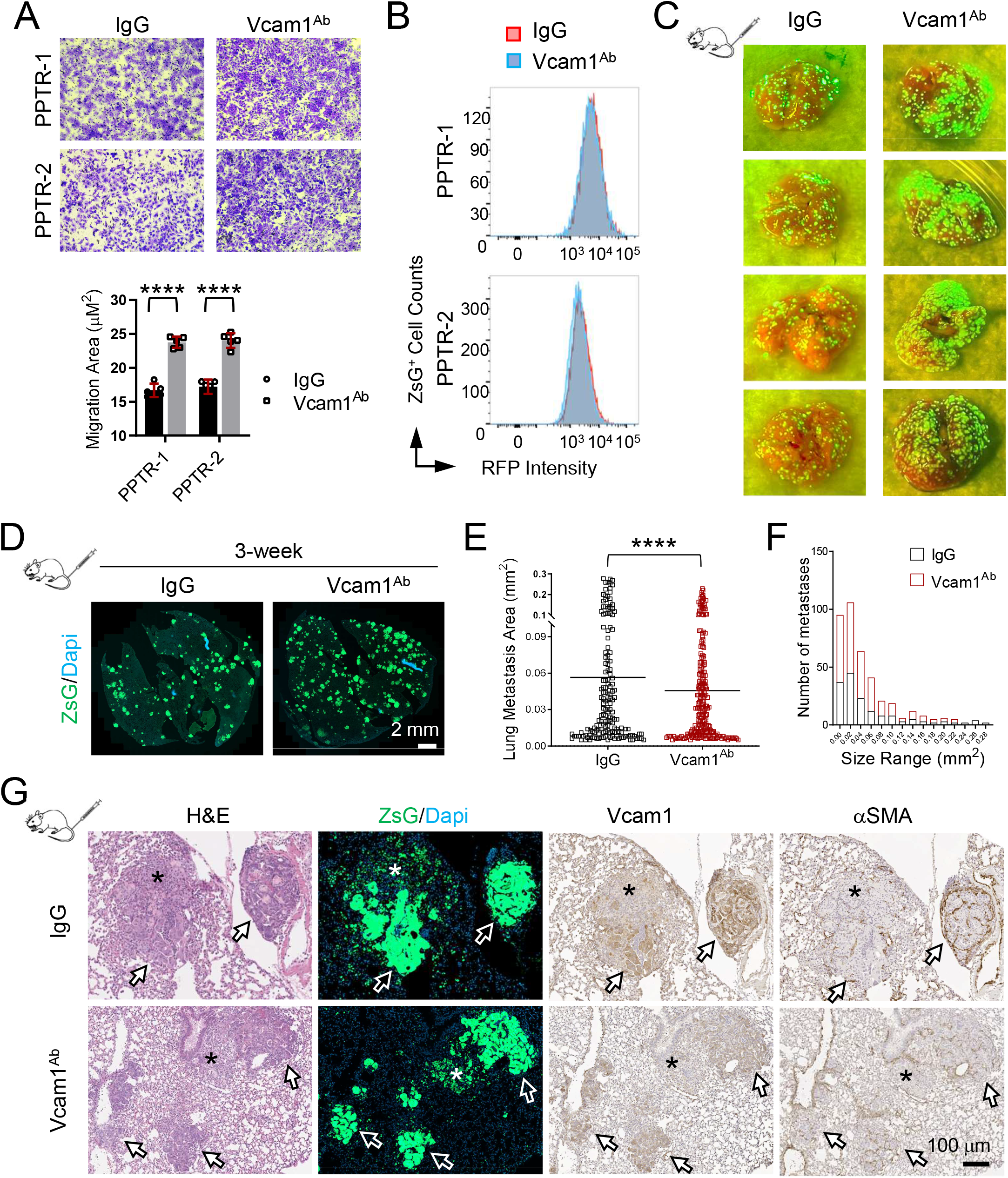
Partial Vcam1 neutralization increases the number of small lung metastases in a TVI model. **(A)** Transwell migration assay and quantification of the PPTR cells treated with IgG or Vcam1^Ab^ showing increased cell migration with Vcam1 blocking. Student *t*-test, *P* value <0.0001. Cells derived from two PPTR tumors were tested. **(B)** Flow cytometry of CellTracker proliferation assay of the PPTR tumor cells treated with IgG or Vcam1^Ab^ found no changes in cell proliferation with Vcam1 blocking. **(C)** Gross ZsG images of the lungs collected from the indicated TVI models three weeks post injection. Vcam1^Ab^ treatment led to more but smaller lung metastases. **(D)** Representative images of the whole-section GFP scans of the lungs from mice treated with IgG or Vcam1^Ab^. **(E)** Quantification of the area of individual lung metastases from the TVI model shows smaller lung metastases in the Vcam1^Ab^ -treated mice. Student *t*-test, *P* value, ****<0.0001. **(F)** Histogram of the size distribution of the lung metastases from the TVI model shows the larger numbers of small lung metastases in the Vcam1^Ab^-treated mice. **(G)** The indicated staining of the lung metastases from the IgG or Vcam1^Ab^ groups. Arrows: Vcam1^+^ tumor cells; asterisk: Vcam1^-^ DTCs. All images share the same 100 μm scale bar.

## DISCUSSION

Using an orthotopic model of metastatic iCCA and multiple coculture models we established in the laboratory, we showed that there was an intriguing MF dynamics during iCCA development. At the early stage of iCCA tumorigenesis, pMFs rapidly accumulate at the tumor border. When tumor progresses, MFs infiltrate into the tumor mass as iMFs while pMFs become enriched at the tumor invasive front. We found iMFs and pMFs play different roles in iCCA progression. iMFs are predominantly growth-promoting as previously reported [3, 4] but with limited effects on tumor cell dissemination. Peritumoral MFs exhibit a surprising suppressive effect on iCCA growth but elicit tumor cell dissemination. Our observations are consistent with the previous observations in HCC patients that activated HSCs and MFs in the peritumoral liver are associated with tumor metastasis and relapse [11, 12], suggesting a role of cell competition in selecting aggressive cancer behaviors [27]. iCCA is known to be highly heterogeneous and plastic [28]. The high rates of tumor relapse in iCCA patients even after curative resection suggests that tumor dissemination is common in these patients. Tumor suppression by pMFs may lead to a selection for aggressive iCCA cells [29].

Based on our findings, we suspect that MF depletion at the early stage of tumorigenesis may target pMFs mostly and dampen the ability of the liver to suppress tumor growth and would lead to accelerated tumor development. However, with continuous increase in MF infiltration as protumorigenic iMFs and weakened ability of pMFs to restrain aggressive tumor growth when tumor progresses, MF depletion in late-stage tumors may in fact an effective strategy to slow tumor development. We suspect that the different timing of MF depletion may account for, at least partially, to the contradicting tumor outcomes in the previously reported CAF-depletion studies [4, 30-33]. A genetic approach capable of temporal-and spatial-specific depletion of MFs, when available, would be the most definitive tool to define the dynamic role of iMFs and pMFs in iCCA development. Single-cell and spatial transcriptomic analyses using our coculture systems and mouse models are also in line to pinpoint the molecular mechanisms behind the differential roles of iMFs and pMFs play in iCCA development.

We show that Vcam1 is upregulated in the tumor cells at the early phase of pMF-iCCA interaction while its level drops when tumor cells begin to disseminate. Although Vcam1 has been previously shown to promote metastasis [22, 23], our tracking of Vcam1 expression in the early-and late-stage metastatic iCCA allograft tumors suggests that its function is beyond pro-or anti-metastatic. Vcam1 facilitates tumor mass expansion in both liver and lung and restrains tumor cells dissemination. Based on our findings in this study, we speculate that the previously reported association between Vcam1 and cancer metastasis may reflect an increased tumor competitiveness with higher Vcam1 level when tumor grows under a growth-suppressive peritumoral TME, rather than a direct promotion of tumor metastasis by Vcam1. Our data argue against the previous proposals to target VCAM1 as a therapeutic strategy to treat metastatic disease [34]. Our results indicate that highly aggressive tumors could potentially benefit from Vcam1 blocking to disseminate more efficiently. A more detailed understanding of the dynamic involvement of Vcam1 in the complex metastasis cascade is needed before we can adequately assess its therapeutic value in metastatic cancer. We acknowledge that this study focuses entirely on the role of Vcam1 in iCCA tumor cells. Vcam1 activity has been reported in other nonmalignant cell types such as endothelial cells and macrophages also plays a critical role in solid tumor metastasis [22, 23]. This further complicates the roles of Vcam1 to iCCA development which warrants more detailed investigation.

For a highly aggressive tumor type like iCCA, the molecular and cellular networks supporting its progression are likely highly plastic and dependent on changes in both intra-and extra-tumoral microenvironment. Our study suggests that iCCA-MF interaction, or tumor -host interaction in a larger picture, is beyond simple pro-or anti-tumorigenic. We theorize that iCCA metastasis is an ongoing competitive process between the attempt of pMFs to block local growth of the tumor and that of tumor cells to break through the suppression. Future studies are in line to identify specific mechanisms underlying this complex tumor-host crosstalk with a goal to simultaneously address local tumor growth and different steps in metastatic cascades.

## ACKNOWLEDGMENTS

We thank Cara Davis-Goodrum, Angie Hammond and other staff members of St. Jude Animal Resource Center Research Services Unit for their technical support on our animal-related studies.

## Abbreviations

iCCA: intrahepatic cholangiocarcinoma
eCCA: extrahepatic cholangiocarcinoma
HCC: hepatocellular carcinoma
MF: myofibroblast
pMF: peritumoral myofibroblast
iMF: intratumoral myofibroblast
CAF: cancer-associated fibroblast
HSC: hepatic stellate cells
pHC: peritumoral hepatocytes
DTC: disseminated tumor cells
PPTR: Prom1^CreERT2^; Pten^flx/flx^; Tp53^flx/flx^; Rosa-ZsGreen
B6: C57BL/6J
ZsG: Rosa-ZsGreen
tdT: Rosa-tdTomato
RFP: Red Fluorescent Protein
GFP: Green Fluorescent Protein
CM: conditioned media
TVI: tail vein injection

## REFERENCES

1 Banales JM, Marin JJG, Lamarca A, Rodrigues PM, Khan SA, Roberts LR et al. Cholangiocarcinoma 2020: the next horizon in mechanisms and management. Nat Rev Gastroenterol Hepatol 2020; 17: 557–588.

2 Rizvi S, Khan SA, Hallemeier CL, Kelley RK, Gores GJ. Cholangiocarcinoma — evolving concepts and therapeutic strategies. Nature Reviews Clinical Oncology 2018; 15: 95–111.

3 Okabe H, Beppu T, Hayashi H, Ishiko T, Masuda T, Otao R et al. Hepatic stellate cells accelerate the malignant behavior of cholangiocarcinoma cells. Ann Surg Oncol 2011; 18: 1175–1184.

4 Affo S, Nair A, Brundu F, Ravichandra A, Bhattacharjee S, Matsuda M et al. Promotion of cholangiocarcinoma growth by diverse cancer-associated fibroblast subpopulations. Cancer cell 2021.

5 Vaquero J, Aoudjehane L, Fouassier L. Cancer-associated fibroblasts in cholangiocarcinoma. Curr Opin Gastroenterol 2020; 36: 63–69.

6 Affo S, Nair A, Brundu F, Ravichandra A, Bhattacharjee S, Matsuda M et al. Promotion of cholangiocarcinoma growth by diverse cancer-associated fibroblast subpopulations. Cancer cell 2021; 39: 866–882 e811.

7 Guedj N, Blaise L, Cauchy F, Albuquerque M, Soubrane O, Paradis V. Prognostic value of desmoplastic stroma in intrahepatic cholangiocarcinoma. Mod Pathol 2021; 34: 408–416.

8 Robinson MW, Harmon C, O’Farrelly C. Liver immunology and its role in inflammation and homeostasis. Cell Mol Immunol 2016; 13: 267–276.

9 Bataller R, Brenner DA. Liver fibrosis. J Clin Invest 2005; 115: 209–218.

10 Yin C, Evason KJ, Asahina K, Stainier DY. Hepatic stellate cells in liver development, regeneration, and cancer. J Clin Invest 2013; 123: 1902–1910.

11 Ji J, Eggert T, Budhu A, Forgues M, Takai A, Dang H et al. Hepatic stellate cell and monocyte interaction contributes to poor prognosis in hepatocellular carcinoma. Hepatology 2015; 62: 481–495.

12 Jiang J, Ye F, Yang X, Zong C, Gao L, Yang Y et al. Peri-tumor associated fibroblasts promote intrahepatic metastasis of hepatocellular carcinoma by recruiting cancer stem cells. Cancer Lett 2017; 404: 19–28.

13 Zhu L, Finkelstein D, Gao C, Shi L, Wang Y, Lopez-Terrada D et al. Multi-organ Mapping of Cancer Risk. Cell 2016; 166: 1132–1146 e1137.

14 Li L, Qian M, Chen IH, Finkelstein D, Onar-Thomas A, Johnson M et al. Acquisition of Cholangiocarcinoma Traits during Advanced Hepatocellular Carcinoma Development in Mice. Am J Pathol 2018; 188: 656–671.

15 Khatamian A, Paull EO, Califano A, Yu J. SJARACNe: a scalable software tool for gene network reverse engineering from big data. Bioinformatics 2019; 35: 2165–2166.

16 Song Y, Kim SH, Kim KM, Choi EK, Kim J, Seo HR. Activated hepatic stellate cells play pivotal roles in hepatocellular carcinoma cell chemoresistance and migration in multicellular tumor spheroids. Sci Rep 2016; 6: 36750.

17 Mederacke I, Dapito DH, Affo S, Uchinami H, Schwabe RF. High-yield and high-purity isolation of hepatic stellate cells from normal and fibrotic mouse livers. Nat Protoc 2015; 10: 305–315.

18 Sha M, Jeong S, Qiu BJ, Tong Y, Xia L, Xu N et al. Isolation of cancer-associated fibroblasts and its promotion to the progression of intrahepatic cholangiocarcinoma. Cancer medicine 2018; 7: 4665–4677.

19 Xu L, Hui AY, Albanis E, Arthur MJ, O’Byrne SM, Blaner WS et al. Human hepatic stellate cell lines, LX-1 and LX-2: new tools for analysis of hepatic fibrosis. Gut 2005; 54: 142–151.

20 Kalluri R. The biology and function of fibroblasts in cancer. Nat Rev Cancer 2016; 16: 582–598.

21 Calon A, Tauriello DV, Batlle E. TGF-beta in CAF-mediated tumor growth and metastasis. Semin Cancer Biol 2014; 25: 15–22.

22 Wieland E, Rodriguez-Vita J, Liebler SS, Mogler C, Moll I, Herberich SE et al. Endothelial Notch1 Activity Facilitates Metastasis. Cancer cell 2017; 31: 355–367.

23 Chen Q, Zhang XH, Massague J. Macrophage binding to receptor VCAM-1 transmits survival signals in breast cancer cells that invade the lungs. Cancer cell 2011; 20: 538–549.

24 Gheldof A, Berx G. Cadherins and epithelial-to-mesenchymal transition. Prog Mol Biol Transl Sci 2013; 116: 317–336.

25 Zhang D, Bi J, Liang Q, Wang S, Zhang L, Han F et al. VCAM1 Promotes Tumor Cell Invasion and Metastasis by Inducing EMT and Transendothelial Migration in Colorectal Cancer. Front Oncol 2020; 10: 1066.

26 Chow A, Huggins M, Ahmed J, Hashimoto D, Lucas D, Kunisaki Y et al. CD169(+) macrophages provide a niche promoting erythropoiesis under homeostasis and stress. Nat Med 2013; 19: 429–436.

27 Parker T, Madan E, Gupta K, Moreno E, Gogna R. Cell Competition Spurs Selection of Aggressive Cancer Cells. Trends Cancer 2020; 6: 732–736.

28 Ma L, Hernandez MO, Zhao Y, Mehta M, Tran B, Kelly M et al. Tumor Cell Biodiversity Drives Microenvironmental Reprogramming in Liver Cancer. Cancer cell 2019; 36: 418–430 e416.

29 Meacham CE, Morrison SJ. Tumour heterogeneity and cancer cell plasticity. Nature 2013; 501: 328–337.

30 Madsen CD. Pancreatic cancer is suppressed by fibroblast-derived collagen I. Cancer cell 2021; 39: 451–453.

31 Chen Y, Kim J, Yang S, Wang H, Wu CJ, Sugimoto H et al. Type I collagen deletion in alphaSMA(+) myofibroblasts augments immune suppression and accelerates progression of pancreatic cancer. Cancer cell 2021; 39: 548–565 e546.

32 Rhim AD, Oberstein PE, Thomas DH, Mirek ET, Palermo CF, Sastra SA et al. Stromal elements act to restrain, rather than support, pancreatic ductal adenocarcinoma. Cancer cell 2014; 25: 735–747.

33 Ozdemir BC, Pentcheva-Hoang T, Carstens JL, Zheng X, Wu CC, Simpson TR et al. Depletion of carcinoma-associated fibroblasts and fibrosis induces immunosuppression and accelerates pancreas cancer with reduced survival. Cancer cell 2014; 25: 719–734.

34 Chen Q, Massague J. Molecular pathways: VCAM-1 as a potential therapeutic target in metastasis. Clin Cancer Res 2012; 18: 5520–5525.

