## Supplemental figures S1-S10 for "Spatiotemporal Dynamics of Vcam1 Regulates Cholangiocarcinoma Mass Expansion and Tumor Dissemination under Growth-suppressive Peritumoral Myofibroblasts"

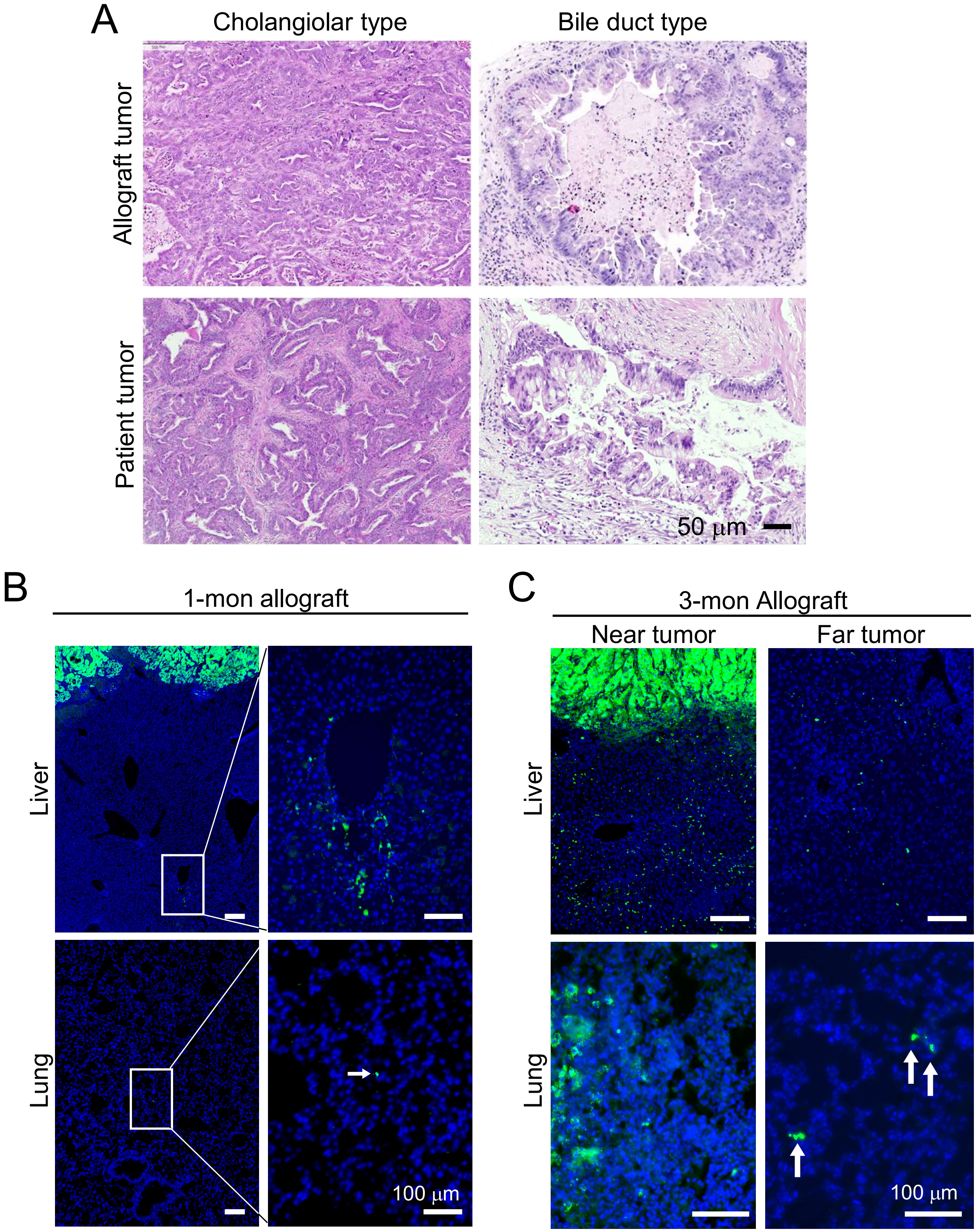


**Supplemental Figure S1. Generation of a metastatic iCCA orthotopic allograft model.**

**(A)** H&E images of the allograft and patient iCCA tumors showing the two types of characteristic histology. All images share the same 50 μm scale bar. **(B, C)** Dissemination of the $ZsG^{+}$ tumor cells in the liver and lung of the 1-month (B) and 3-month (C) iCCA allograft tumors. Arrows: DTCs in the lung. All scale bars are 100 μm.

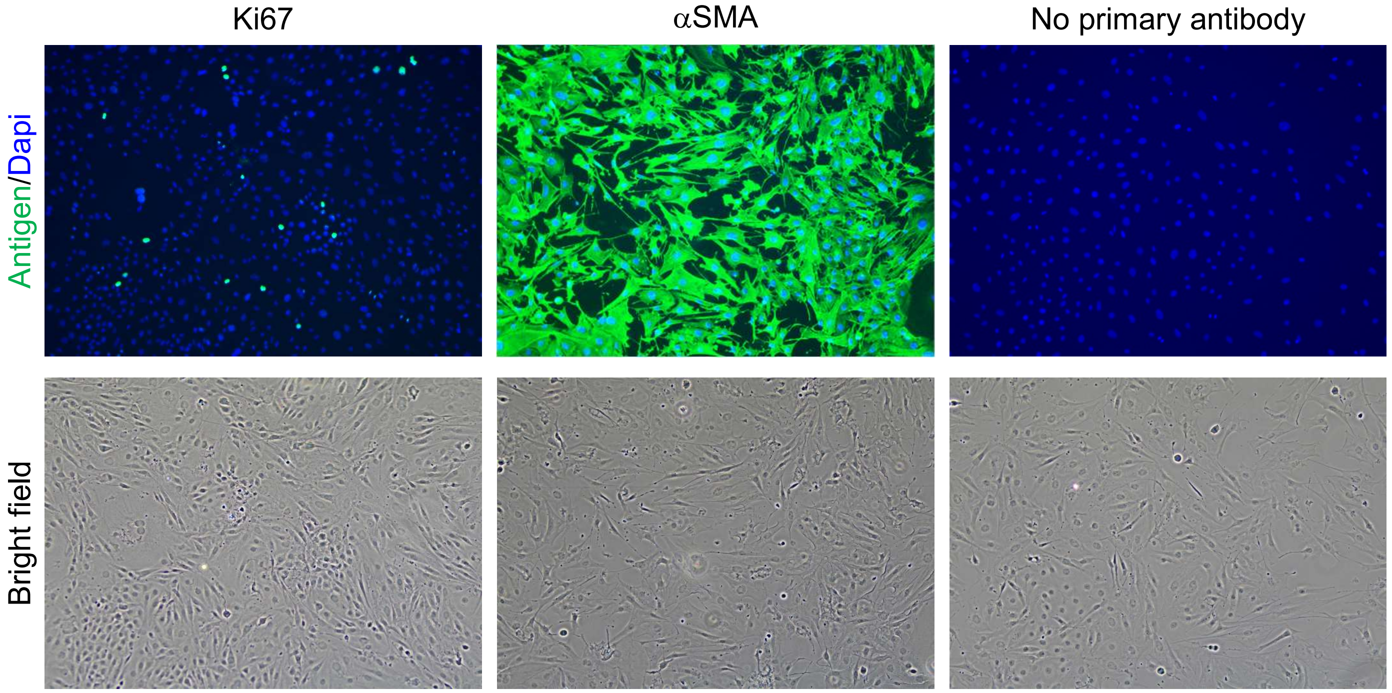


**Supplemental Figure S2. Activation of mouse primary HSCs in 2D culture.**

Ki67 and αSMA immunofluorescence staining of the mouse primary HSCs after three passages on 2D plastic plates, showing their active proliferation and strong expression of the MF marker αSMA.

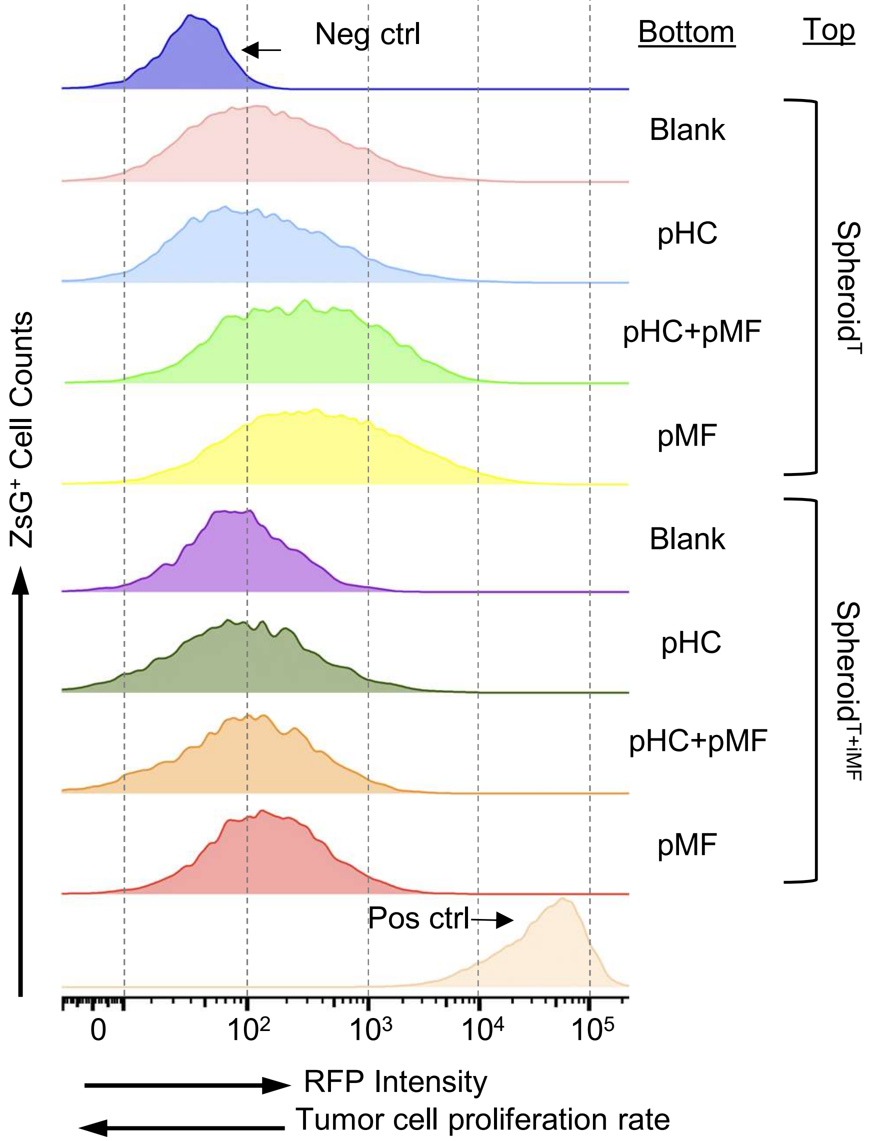


**Supplemental Figure S3. Peritumoral MFs suppress, and intratumoral MFs promote, iCCA growth in vitro.**

RFP intensity histogram of the tumor cells in the indicated coculture conditions in Figure 2C. Higher RFP intensity indicates slower cell proliferation.

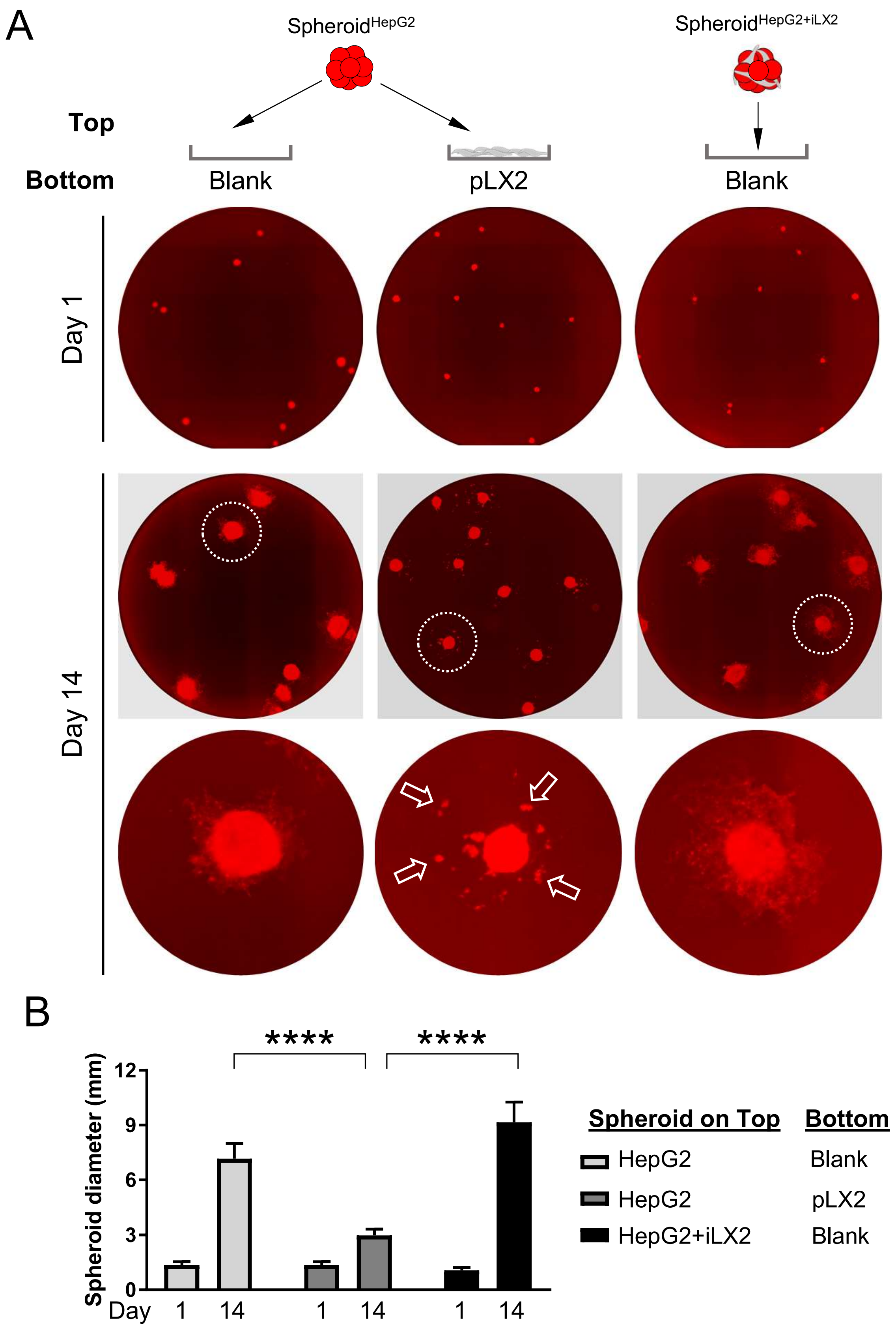


**Supplemental Figure S4. The impact of pMF and iMF on HepG2 cells in 2.5D coculture.**

**(A)** Day 1 and Day 14 RFP image of HepG2 cells cocultured with a human HSC cell line LX2 seeded according to the diagram illustrated on the top. Arrows: disseminated HepG2 cells. **(B)** Quantification of the diameter of the spheroids in he indicated culture on Day 1 and Day 14. Multiple *t* test, *P* value **** <0.0001.

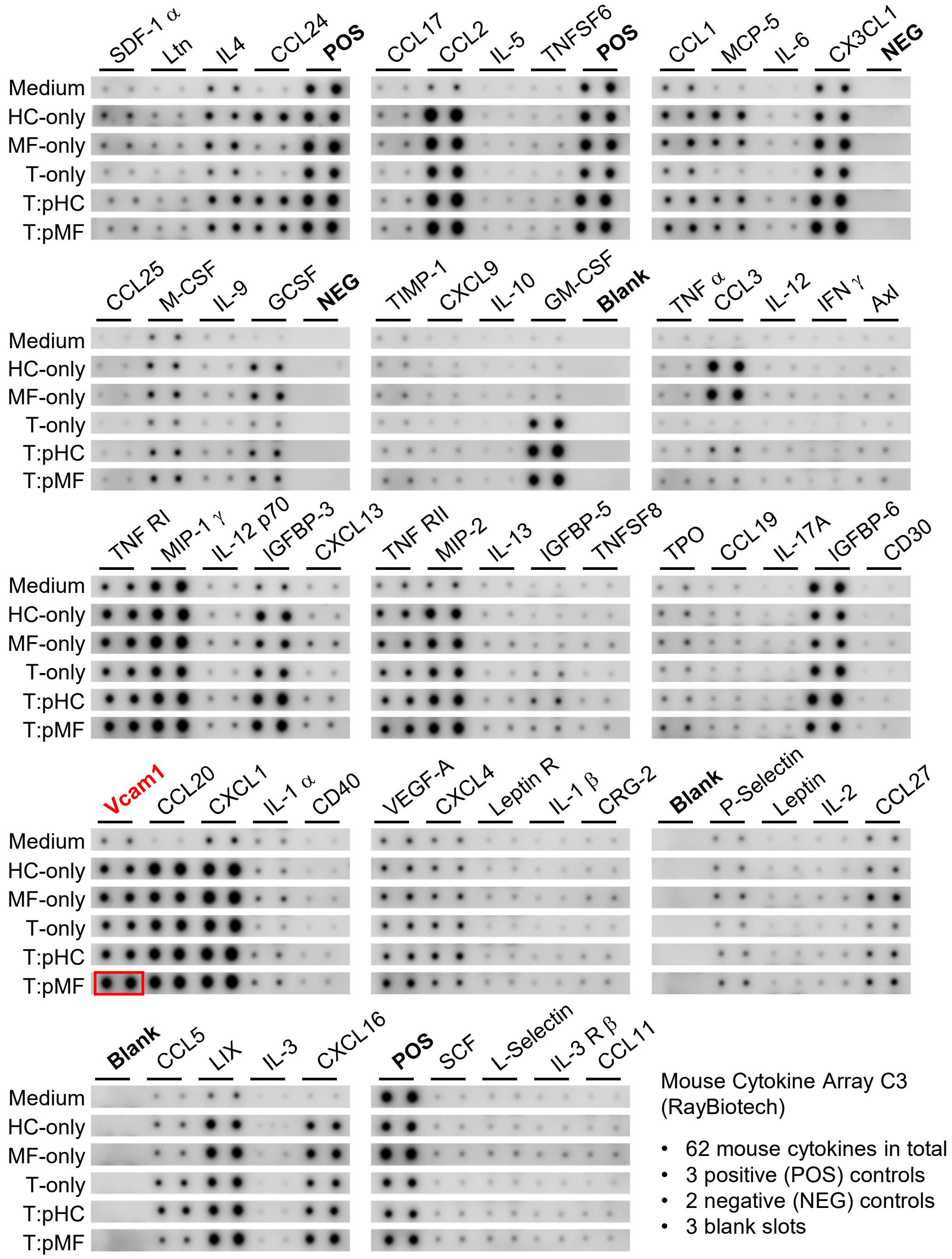


**Supplemental Figure S5. Mouse cytokine array assay identified Vcam1 upregulation in tumor-pMF coculture.**

Individual rows of the cytokine array membranes shown in **Figure 6A** were cropped and aligned for a better visual comparison of the cytokine levels between the five indicated culture conditions. Red boxes: Vcam1 was the only cytokine that showed evident and specific increase specifically in T:pMF coculture.

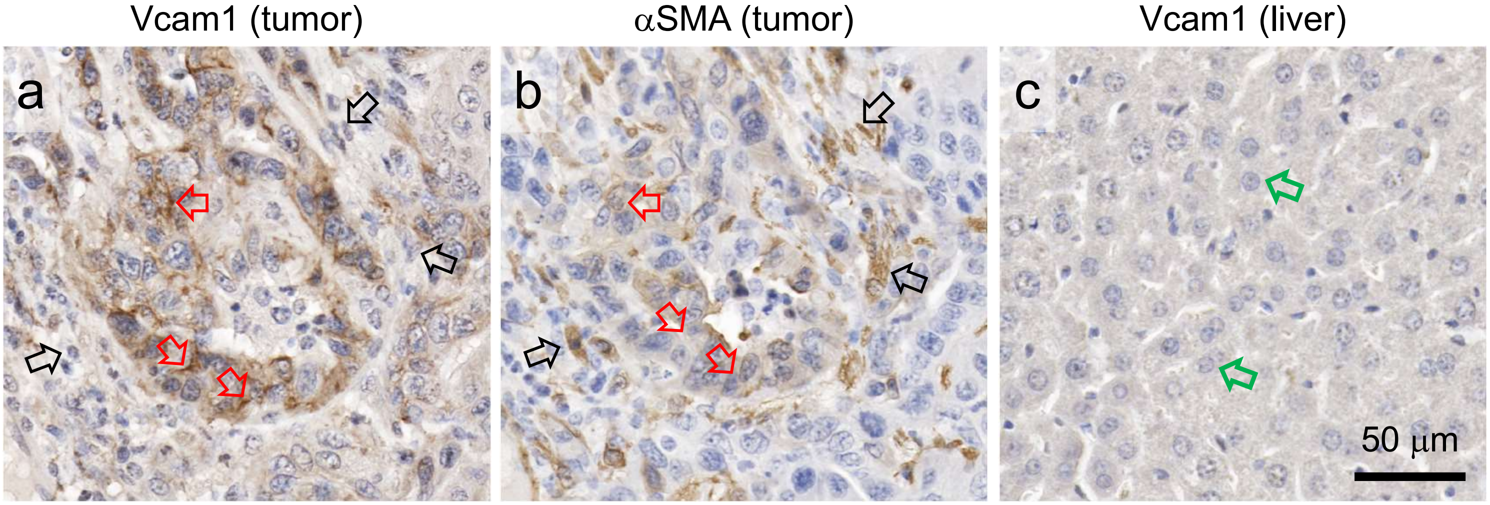


**Supplemental Figure S6. Vcam1 is expressed by tumor cells not MFs.**

IHC staining of **(a)** Vcam1 in the tumor area, **(b)** aSMA in the tumor area, and **(c)** Vcam1 in the non-tumor area with predominantly HCs. Red arrows: cell membrane between tumor cells; black arrows: MFs; green arrows: HCs.

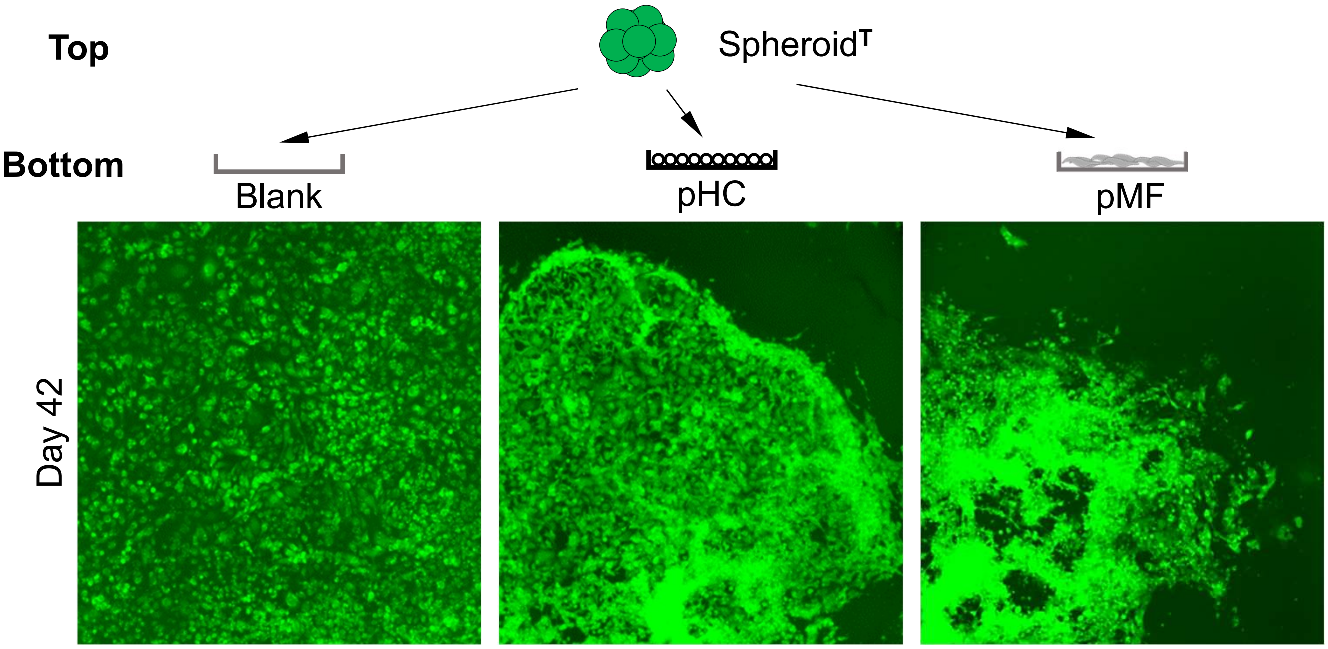


**Supplemental Figure S7. Long-term PPTR 2.5D culture.**

Day 42 ZsG fluorescence images of the PPTR Spheroid$^T$ cultured without a bottom layer (blank), on top of pHCs, or on top of pMFs.

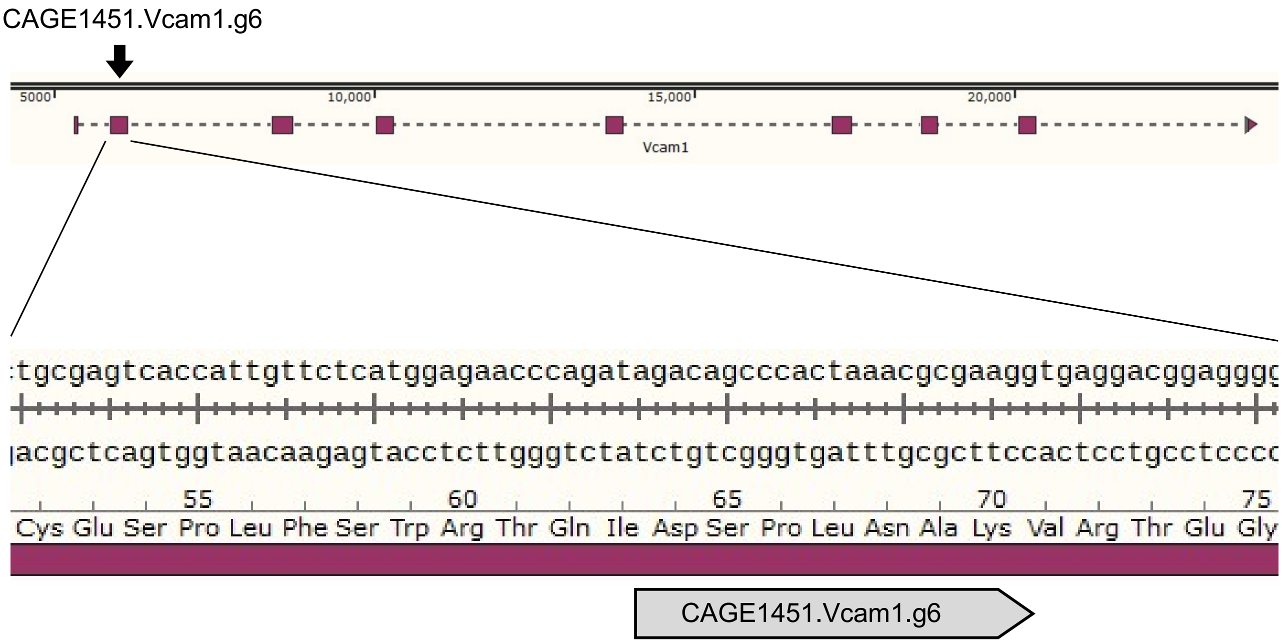


**Supplemental Figure S8. CRISPR/Cas9 knockout targeting of mouse *Vcam1* gene.**

Guide RNA CAGE1451.Vcam1.g6 was used to target mouse *Vcam1* gene.

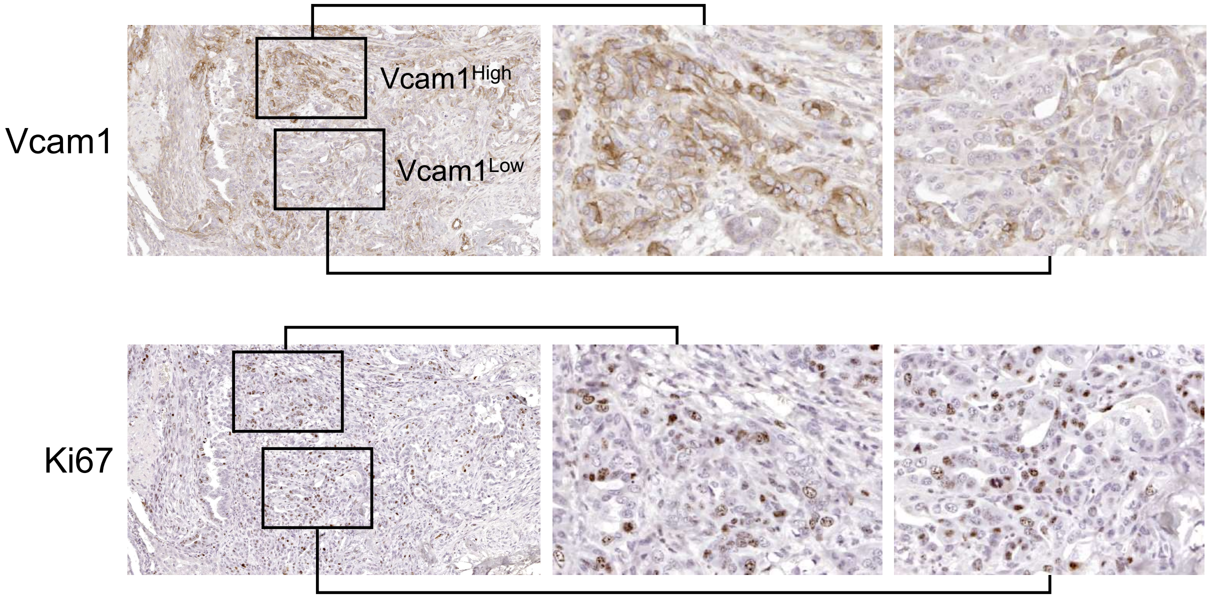


**Supplemental Figure S9. Vcam1 is not associated with PPTR tumor cell proliferation in vivo.**

Vcam1 and Ki67 IHC on serial sections of a 3-month iCCA allograft tumor. No association was found between Vcam1 expression and Ki67 positivity.

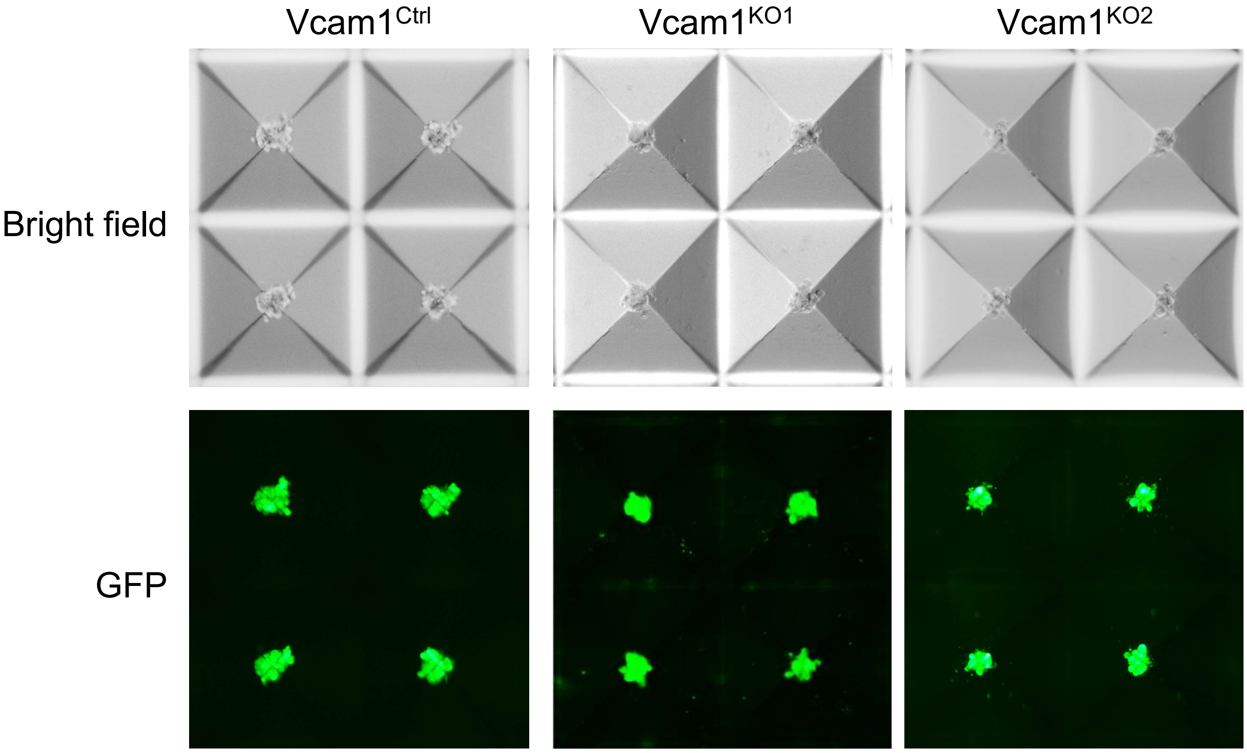


**Supplemental Figure S10.** ***Vcam1$^{KO}$*** **cells were able to form spheroids in AggreWell within 48 hours.**
