## Supplemental methods for "Spatiotemporal Dynamics of Vcam1 Regulates Cholangiocarcinoma Mass Expansion and Tumor Dissemination under Growth-suppressive Peritumoral Myofibroblasts"

**SupplementaL MaterialS**

**Mice**

Two-month-old male and female C57BL/6J (B6) (Strain # 000664, The Jackson Laboratory, Bar Harbor, ME, USA) and *Crl:CD1-Foxn1^nu/nu^* (CD-1 nude) mice (Strain Code 086, Charles River, Wilmington, MA, USA) were used for iCCA orthotopic transplantation and tail vein injection. For iCCA orthotopic transplantation, PPTR tumor cells were surgically injected into the mouse liver at 1×10^5^/mouse in 4 μl cold growth factor-reduced matrigel (Cat. No. 354230, Corning Inc, Corning, NY). For tail vein injection (TVI) lung metastasis model, tumor cells were injected at 1×10^6^/mouse to CD-1 nude mice in 100 μl PBS via TVI. *B6.129(Cg)-Gt(ROSA)26Sor^tm4(ACTB-tdTomato,-EGFP)Luo^/J* (Strain No. 007676, The Jackson Laboratory, Bar Harbor, ME, USA) mice were used to isolated hepatocytes according to standard protocol. Equal numbers of males and females were randomly selected and included in each in vivo assays. All mice were maintained in the Animal Resource Center at St. Jude Children’s Research Hospital (St. Jude). Animal protocols were approved by the St. Jude Animal Care and Use Committee.

**Cell Culture**

1. **PPTR tumor cell culture**

PPTR cells were established from *Prom1^CreERT2^; Pten^flx/flx^; Tp53^flx/flx^; Rosa-ZsGreen* (PPTR) liver cancer organoids [1] and adapted to 2D culture in DMEM (Cat. No. 15-013-CV, Corning Inc, ) supplemented with 10% fetal bovine serum (FBS) (Cat. No. 10082147, Gibco, Waltham, MA) and antibiotics. Primary mouse HSCs were purchased (Cat. No. M5300-57, ScienCell Research Laboratories, San Diego, CA, USA) and cultured according to the manufacturer’s instruction. All iCCA-liver coculture was maintained in a 1:1 mixture of in Stellate Cell Medium (Cat. No. 5301, ScienCell) and the DMEM-based tumor culture medium (Corning). AggreWell 400 culture plates (Cat. No. #34415, STEMCELL, Cambridge, MA) were used to generate tumor spheroids. Two-chamber culture insert and µ-slide 8-well Grid-500 culture slides (Cat. No. 501149022, ibidi, Gräfelfing, Germany) were used in 2D coculture. All coculture-related assays were performed in at least three biological replicates.

2. **Primary liver cell isolation**

Liver cells were isolated from two-month-old *mTmG* mouse liver using the standard two-step collagenase perfusion method [2, 3]. Briefly, mice were anesthetized with isoflurane, the liver was perfused with 30 ml of prewarmed EGTA solution (0.5 mM) through the inferior vena cava for 10 min, and then perfused with 30 ml of prewarmed protease solution (14 mg/mouse; Cat. No. P5147-5G, MilliporeSigma, Burlington, MA) followed by 30 ml of collagenase solution (3.7 U/mouse; Cat. No. 11088882001, Roche, Basel, Switzerland) for 10 min each. The liver was then removed and transferred to a sterile Petri dish and gently minced with forceps. The liver was further digested with the protease/collagenase solution with 1% DNase I (Cat. No. 10104159001, Roche) for 25 min at 40 ℃. Cells were filtered by a 70-μm filter and centrifugated at 50 × *g* for 3 min to pellet HCs. HCs were then washed twice in cold Advanced DMEM/F12 (Cat. No. 12634028, Thermo Fisher Scientific, Waltham, MA, USA) and subjected to culture.

3. **Hepatocyte 2D culture**

Freshly isolated HCs were subjected to 3D organoids culture as previously reported for three passages.[1] HC organoids were then dissociated and transferred to 60mm culture dishes precoated with 1% GFR matrigel (Corning) and culture in 2D condition using the same organoid culture medium.

4. **Mouse hepatic stellate cell culture**

Primary mouse HSCs were purchased (Cat. No. M5300-57, ScienCell Research Laboratories, San Diego, CA, USA) and cultured in Stellate Cell Medium (Cat. No. 5301, ScienCell) supplemented with 1% stellate cell growth supplement (Cat. No. 5352, ScienCell), 2% FBS (Gibco), and 1% Penicillin/Streptomycin (Cat. No. TMS-AB2-C, MilliporeSigma) Solution.

5. **Spheroid culture**

AggreWell 400 (AW400) culture plates (STEMCELL) were used to generate tumor spheroids. Spheroid^T^ was generated using 200 tumor cells/microwell and Spheroid^T+iMF^ was generated using 100 tumor cells+100 MFs/microwell, allowed to aggregate for 48 hours and then transferred to 2.5D cultures. DMEM (Corning) supplemented with 10% fetal bovine serum (FBS) (Gibco) and antibiotics was used to culture tumor spheroids. The cholangiocyte organoid culture medium was prepared by supplementing advanced DMEM/F12 (Thermo Fisher) with 50% conditioned medium from L-WRN cells (Cat. No. CRL-3276™, ATCC, Manassas, VA), 10 mM HEPES (Cat. No. H3375-25G, MilliporeSigma), 1% GlutaMax(Cat. No. 35050061, Thermo Fisher Scientific), 1% Penicillin-Streptomycin (MilliporeSigma), 2% B27 (Cat. No. 17504001, Thermo Fisher Scientific), 1% N2 (Cat. No. 17502001, Thermo Fisher Scientific), 3 μm CHIR 99021(Cat. No. 4423, Bio-Techne Corporation, Minneapolis, MN), 1.25 mM N-acetylcysteine (Cat. No. A0737, MilliporeSigma), 10 mM Nicotinamide (Cat. No. N0636, MilliporeSigma), 10 nM recombinant gastrin (Cat. No. SCP0152, MilliporeSigma), 50 ng/ml EGF (Cat. No. 500-P45, Peprotech, Cranbury, NJ), 50 ng/ml FGF7 (Cat. No. 100-19, Peprotech), 50 ng/ml FGF10 (Cat. No. 100-26, Peprotech), 25 ng/ml HGF (Cat. No. 100-39, Peprotech), 1 μM A83-01 (Cat. No. 2939, Bio-Techne Corporation), 10 μM Rho Inhibitor γ-27632 (Cat. No. 72308, STEMCELL), and 1% growth factor-reduced matrigel (Corning).

6. **iCCA-liver cocultures**

All iCCA-liver coculture was maintained in a 1:1 mixture of in Stellate Cell Medium (Cat. No. 5301, ScienCell) and the DMEM-based (Corning) tumor culture medium.

(1) **Tumor-HC-MF 2.5D coculture (Figure 2C and Figure 3A)**: Freshly isolated *mTmG* HCs were mixed with primary mouse MFs at a ratio of HC:MF = 10:0 (HC-only), 8:2, 5:5, 2:8, or 0:10 (MF-only), seeded in 96-well flat-bottom microplates precoated with 1% GFR matrigel at 2×10^4^ total cells/well, and cultured in a 1:1 mixture of tumor culture medium and Stellate Cell Medium for 24 hr. PPTR tumor spheroids were generated as in (3) above and added at 2-5 spheroids/well. The cocultures were maintained up to 42 days and the medium was refreshed every 3-4 days. The ZsG fluorescence images were capture for the first four days, the diameters of three spheroids from three biological replicates were in each condition measured in Image J and plotted in GraphPad Prism 7.
(2) **Tumor-MF 2.5D coculture** (**Figure 6E**): MFs were seeded at 2×10^4^ cells/well in the 96-well microplates and tumor spheroids were seeded on top after 24 hours. The same culture medium as in (1) was used.
(3) **Tumor-HC-MF 2D coculture (Figure 3C)**: Two-chamber culture insert was placed in the µ-slide 8-well Grid-500 culture slides (ibidi) and the culture surface was precoated with 1% GFR matrigel (Corning) at 37°C for 24 hr. PPTR tumor cells, MFs and 2D-cultured TdT^+^ HCs were seeded in various combinations as indicated in Figure 4C. The total number of cells in each chamber was kept at 4.25×10^4^ cells and the mixing ration of T:MF or HC:MF was 4:1. MFs were labeled with CellTracker^TM^ Red CMTPX (Cat. No. C34552, Invitrogen, Carlsbad, CA) in the conditions TdT^+^ HCs were not used. The cocultures were maintained for 14 days and the culture medium was refreshed every 3–4 days.

**iCCA Transplantation Models**

Based on in vitro data, we expected to detect a large standardized effect size. We relaxed the type I error rate to 15% with 80% power and an expected effect size of 0.9. For the three types of in vivo tests, a total of 12 mice were needed, i.e., 4 mice for each subgroup.

1. **iCCA orthotopic transplantation model**

PPTR tumor cells were surgically injected into the liver of two-month-old male and female CD-1 nude mice or B6 mice. Briefly, a midline longitudinal abdominal incision was made to expose liver left lobe and tumor cells were injected at 1×10^5^/mouse in 4 μl cold growth factor-reduced (GFR) matrigel (Corning) using a Hamilton syringe and needle (Model No. 7105, Hamilton Company, Bonaduz, Switzerland). Survival curves and median survival of the orthotopic allograft models were determined by the Kaplan–Meier method in GraphPad Prism 7.

2. **iCCA tail vein injection (TVI) lung metastasis model**

PPTR tumor cells were injected into two-month-old male and female CD-1 nude mice at 1×10^6^/mouse in 100 μl PBS via TVI. The lung was examined three weeks post injection.

**CellTracker^TM^ Cell Proliferation Assay**

PPTR cells were labeled by 5 μM CellTracker^TM^ Red CMTPX dye (Invitrogen) in Advanced DMEM/F12(Thermo Fisher) without FBS for 1 hour, then washed with Advanced DMEM/F12 (Thermo Fisher). 200 tumor cells/microwell and 100 PPTR + 100 MF cells/microwell in AW400 plates to aggregate to the spheroids. The culture medium was a mixture of 50% cholangiocyte organoid culture medium and 50% Stellate Cell Medium (Cat. No. 5301, ScienCell). HCs and MFs at a ratio of HC-only, 5:5, MF-only, and blank cultured in 96-well flat-bottom microplates. The culture medium was a 1:1 mixture of in Stellate Cell Medium and the DMEM-based tumor culture medium. After 2 days, transferred around 2-3 spheroids/well into the 96-well plates. After 4 days culture, the cells have been collected and washed with 1x DPBS (Cat. No. 21-031-CV, Corning). Cells were resuspended with 100 μl 2% FBS (Gibco) in DPBS (Corning) solution and read the RFP using flow cytometry.

**Cytokine array assay**

Conditioned media (CM) were collected from cell cultures after four days. The levels of secreted cytokines in the CM were measured using Mouse Cytokine Antibody Array 3 (Cat. No. AAM-CYT-3-8, RayBiotech, GA) according to the manufacturer's instructions. Briefly, 1 ml of undiluted CM were incubated with the Mouse Cytokine Antibody Array C3 membranes overnight at 4°C. After washing, the membranes were incubated with the Biotinylated Antibody Cocktail and subsequently with HRP-Streptavidin, 2 hours at room temperature for each. After washing, the cytokine signals were visualized via the LI-COR chemiluminescence imaging system. The signals were quantified using Image Studio Lite Ver5.2 (LI-COR) and normalized to the positive controls on the membrane.

**Quantitative RT-PCR**

The total RNA in cell pellets were extracted using RNeasy® Mini Kit (Qiagen, Hilden, Germany). SuperScript® III First-Strand Synthesis SuperMix for qRT-PCR (Invitrogen) was used for cDNA synthesis from 500 ng total RNA. FastStart Universal SYBR® Green Master (ROX) (Roche) was used to perform the quantitative PCR assay. The results were analyzed using 2^-ΔΔCt^ method, with ATCB (β-actin) as the internal reference gene. Quantitative RT-PCR was performed using three batches of independent cell cultures and the result was plotted using GraphPad Prism. The PCR primers were as follows: Vcam1: GCCCACTAAACGCGAAGGT (Forward), ATGGTCAGAACGGACTTGGAC (Reverse); β-actin: GTTGTCGACGACCAGCG (Forward), GCACAGAGCCTCGCCTT (Reverse).

**Cell Migration Assay**

Cell migration capacity was measured using polycarbonate membrane inserts (Cat. No. CBA-100-COL, Cell BioLabs, San Diego, CA) in 24-well plates. In brief, PPTR cells were seeded in 60mm dish and treated with 5 μM Vcam1^Ab^ (Cat. No. #BE0027, Bioxcell, Lebanon, NH) or IgG (Cat. No. Mab006, R&D System, Minneapolis, MN) for 4 days. PPTR cell suspension was then added to the upper chamber (3 × 10^4^ cells in 300 µl blank medium with 5 μM of Vcam1 antibody or IgG, respectively), and 500 µl of media with 10% FBS (Gibco) was added to the lower chamber. After incubation at 37 °C under 5% CO_2_ for 24 h, the cells on the upper surface were wiped with cotton swabs, while the cells that migrated through the filter pores were stained with cell stain solution for 10 min. The stained cells were observed and photographed under an inverted microscope. Cell area was measured in five randomly selected fields using Image J and plotted in GraphPad Prism 7.

**Immunohistochemistry and immunofluorescence**

Paraffin sections (4 µm) of liver spheroids and tissues were prepared by HistoWiz Inc. (Brooklyn, NY, USA) and analyzed by direct fluorescence microscopy, H&E staining, and IHC. Primary HSCs cultured on glass chamber were subject to standard immunofluorescence. Antibodies used included anti-Ki67 (Cat. No. ab16667, Abcam, Cambridge, MA, 1:200); anti-αSMA (Cat. No. ab124964, Abcam, 1:1000); anti-Vcam1 (Cat. No. ab134047, Abcam, 1:1000); and anti-E-Cadherin (Cat. No. ab231303, Abcam, 1:1000).

**Microscopy, Image-based Quantification and Statistical Analysis**

All cocultures were monitored daily by using an ECLIPSE Ts2R fluorescence microscope (Nikon, Minato City, Tokyo, Japan) and Lionheart FX Automated Microscope (Winooski, VT, USA). The ZsG+ tumor cell area was measured by Image J and plotted in GraphPad Prism 7. Two-tailed student t test was performed in GraphPad Prism 7 to compare two independent pairs of groups. Two-way ANOVA was performed when two or more groups were compared. P value ≤0.05 was considered statistically significant. All the quantifications below were performed on three biological replicates in each group.

1. **Area of iMF vs. pMF in patient and mouse iCCA:** three regions with the heaviest iMF or pMF accumulation were selected from the patient and mouse iCCA tumors and the area occupied by αSMA staining signal was measured in Image J and plotted in GraphPad Prism 7. Student *t* test was performed for comparison.
2. **Tumor cell proliferation in vivo:** PPTR cells in the orthotopic tumors retained their direct ZsG fluorescence after formalin fixation and paraffin embedding. To quantify Ki67^+^ tumor cells, tumor sections were deparaffinization and scanned for ZsG fluorescence prior to Ki67 IHC. After the completion of IHC, 200× images were captured from three independent tumor areas for each group. The number of nuclei of ZsG^+^/Ki67^+^ cells were manually counted and plotted in GraphPad Prism 7. Student *t* test was performed for comparison.
3. **Tumor spheroid expansion:** The ZsG fluorescence images were capture for the first four days from each condition and the long and short diameters of three spheroids from each condition were measured in Image J and plotted in GraphPad Prism 7. Two-way ANOVA was performed for comparison.
4. **Tumor cell invasion in 2.5D coculture:** tumor cell invasive processes were measured in length using Image J and plotted in GraphPad Prism 7. Two-way ANOVA was performed for comparison.
5. **Tumor cell expansion in the two-chamber coculture:** ZsG/bright-fiend images of the cocultures from Day 0 and 14 were aligned in Photoshop using the 500-μm engraved grid on the culture slides. The movement distance of the tumor border was measured from three independent sets of experiments in Image J and plotted in GraphPad Prism 7. Student *t* test was performed for comparison.
6. **Lung metastasis area measurement:** Whole lung ZsG fluorescence images were captured by AxioScan Z.1 Whole Slide Scanner (Zeiss, Oberkochen, Germany). The number of lung metastases and the area of individual metastasis were measured in Image J and plotted in GraphPad Prism 7. Student *t* test was performed for comparison. Five lungs each from the IgG vs. Vcam1^Ab^ group or Vcam1^Ctrl^ vs. Vcam1^KO^ were compared.
7. **Cell migration:** Cell area was measured in five randomly selected fields using Image J and plotted in GraphPad Prism 7. Student *t* test was performed for comparison.

**Total mRNA sequencing and analysis**

Total RNA library of the *Vcam1^Ctrl^* and *Vcam1^KO^* PPTR cells was constructed using Illumina TrueSeq stranded mRNA library prep kit and and paired-end 100-cycle sequencing was performed on Illumina NovaSeq sequencers per the manufacturer’s directions (Illumina, San Diego, CA). Reads were then counted by pseudoalignment method kallisto (v0.46.1, parameters “quant --rf-stranded --plaintext ---bias”) based on mouse genome mm10 (Gencode GRCm38 vM24) transcripts annotations [4]. All samples achieved > 50 million fragments and > 90% mapping rate. After trimmed mean of M-values normalization (edgeR v3.34.0) [5], Voom (limma v3.48.3) [6] was used to identify differential expressed genes. Enrichr server were used for pathway enrichment analysis [7]. GSEA(v4.0.3) [8] Prerank mode were ran with MSigDB(v7.4) [9] against the gene rank by log2 fold change.

**Gene regulatory network analysis**

A scalable software was used for gene regulatory network reverse-engineering from big data, SJARACNe (v-0.1.0) [10], to reconstruct context-dependent signaling interactomes of Vcam1. The adaptive partitioning algorithm was used for mutual information estimation. The upstream first neighbors of Vcam1 were extracted and considered as the regulators in each context. A hypergeometric distribution method was applied for the gene set enrichment analysis using the “funcEnrich.Fisher” function from the R package NetBID (v-2.0.2) [11]. Only the HALLMARK and KEGG gene sets from the MSigDB database (v-6.1) [8] were used. The *P* values of the gene set enrichment analysis of both PPTR and TCGA datasets were combined using the Stouffer method embedded in the “combinePvalVector” function from NetBID. The visualization was completed by ggplot2 (v-3.3.4, Wickham H (2016). ggplot2: Elegant Graphics for Data Analysis. Springer-Verlag New York. ISBN 978-3-319-24277-4, <https://ggplot2.tidyverse.org>).

**References**

1	Li L, Qian M, Chen IH, Finkelstein D, Onar-Thomas A, Johnson M *et al*. Acquisition of Cholangiocarcinoma Traits during Advanced Hepatocellular Carcinoma Development in Mice. *Am J Pathol* 2018; 188: 656-671.

2	Berry MN, Friend DS. High-yield preparation of isolated rat liver parenchymal cells: a biochemical and fine structural study. *J Cell Biol* 1969; 43: 506-520.

3	Li WC, Ralphs KL, Tosh D. Isolation and culture of adult mouse hepatocytes. *Methods Mol Biol* 2010; 633: 185-196.

4	Harrow J, Frankish A, Gonzalez JM, Tapanari E, Diekhans M, Kokocinski F *et al*. GENCODE: the reference human genome annotation for The ENCODE Project. *Genome Res* 2012; 22: 1760-1774.

5	Robinson MD, McCarthy DJ, Smyth GK. edgeR: a Bioconductor package for differential expression analysis of digital gene expression data. *Bioinformatics* 2010; 26: 139-140.

6	Law CW, Chen Y, Shi W, Smyth GK. voom: Precision weights unlock linear model analysis tools for RNA-seq read counts. *Genome Biol* 2014; 15: R29.

7	Kuleshov MV, Jones MR, Rouillard AD, Fernandez NF, Duan Q, Wang Z *et al*. Enrichr: a comprehensive gene set enrichment analysis web server 2016 update. *Nucleic Acids Res* 2016; 44: W90-97.

8	Subramanian A, Tamayo P, Mootha VK, Mukherjee S, Ebert BL, Gillette MA *et al*. Gene set enrichment analysis: a knowledge-based approach for interpreting genome-wide expression profiles. *Proc Natl Acad Sci U S A* 2005; 102: 15545-15550.

9	Liberzon A, Birger C, Thorvaldsdottir H, Ghandi M, Mesirov JP, Tamayo P. The Molecular Signatures Database (MSigDB) hallmark gene set collection. *Cell Syst* 2015; 1: 417-425.

10	Khatamian A, Paull EO, Califano A, Yu J. SJARACNe: a scalable software tool for gene network reverse engineering from big data. *Bioinformatics* 2019; 35: 2165-2166.

11	Du X, Wen J, Wang Y, Karmaus PWF, Khatamian A, Tan H *et al*. Hippo/Mst signalling couples metabolic state and immune function of CD8alpha(+) dendritic cells. *Nature* 2018; 558: 141-145.
